# A survey of hypothalamic phenotypes identifies molecular and behavioral consequences of MYT1L haploinsufficiency in male and female mice

**DOI:** 10.1101/2024.11.25.625294

**Authors:** Susan E. Maloney, Katherine B. McCullough, Sneha M. Chaturvedi, Din Selmanovic, Rebecca Chase, Jiayang Chen, Doris Wu, Jorge L. Granadillo, Kristen L. Kroll, Joseph D. Dougherty

**Affiliations:** Department of Psychiatry, Washington University School of Medicine, St. Louis, MO, USA; Intellectual and Developmental Disabilities Research Center, Washington University School of Medicine, St. Louis, MO, USA; Department of Genetics, Washington University School of Medicine, St. Louis, MO, USA; Department of Pediatrics, Division of Genetics and Genomic Medicine, Washington University School of Medicine, St. Louis, MO, USA; Department of Developmental Biology, Washington University School of Medicine, St. Louis, MO, USA

**Keywords:** MYT1L, mouse model, haploinsufficiency, hypothalamus, feeding, gene expression, arginine vasopressin, oxytocin, maternal care, aggression

## Abstract

The transcription factor MYT1L supports proper neuronal differentiation and maturation during brain development. MYT1L haploinsufficiency results in a neurodevelopmental disorder characterized by intellectual disability, developmental delay, autism, behavioral disruptions, aggression, obesity and epilepsy. While MYT1L is expressed throughout the brain, how it supports proper neuronal function in distinct regions has not been assessed. Some features of MYT1L Neurodevelopmental Syndrome suggest disruption of hypothalamic function, such as obesity and endocrine issues, and previous research showed changes in hypothalamic neuropeptide expression following knockdown in zebrafish. Here, we leveraged our heterozygous *Myt1l* mutant, previously shown to recapitulate aspects of the human syndrome such as hyperactivity, social challenges, and obesity, to examine the impact of MYT1L loss on hypothalamic function. Examining the molecular profile of the MYT1L haploinsufficient hypothalamus revealed a similar scale of disruption to previously studied brain regions, yet with region-specific roles for MYT1L, including regulation of neuropeptide systems. Alterations in oxytocin and arginine vasopressin cell numbers were also found. Behaviors studied included maternal care, social group hierarchies, and aggression, all of which were unchanged. Feeding and metabolic markers were also largely unchanged in MYT1L haploinsufficient mice, yet an interaction was observed between diet and MYT1L genotype on weight gain. Our findings here suggest that gross endocrine function was not altered by MYT1L haploinsufficiency, and that key sex-specific behaviors related to proper hypothalamic function remain intact. Further study is needed to understand the functional impact of the altered hypothalamic molecular profile and changes in neuropeptide cell numbers that result from MYT1L haploinsufficiency.

## Introduction

MYT1L (Myelin Transcription Factor 1 Like) is a zinc finger transcription factor that is highly expressed in the developing brain and plays an important role in neuronal maturation (Yen et al., 2024). MYT1L promotes neuronal differentiation while inhibiting non-neuronal gene expression, acting both as a repressor and an activator (Chen et al., 2022). Heterozygous loss-of-function mutations in *MYT1L* are known to cause a rare autosomal dominant disorder in humans characterized by intellectual disability of varying degree, global developmental delay, autism, aggression and behavioral disorders, obesity, and epilepsy (Coursimault et al., 2022). Mice with heterozygous constitutive *Myt1l* mutations have been shown to recapitulate aspects of the human phenotype, presenting with similar features such as obesity, microcephaly, social impairments, hyperactivity, and altered motor functions with reduced strength suggestive of hypotonia (Chen et al., 2021; Kim et al., 2022; Wöhr et al., 2022). MYT1L is expressed in neurons of all brain regions; however, the effect of MYT1L haploinsufficiency across all regions has not been assessed.

There is some suggestion that a portion of the MYT1L Neurodevelopmental Syndrome phenotype may be related to dysfunction of the hypothalamus. Notably, the patients tend to be obese, suggesting feeding or metabolic anomalies, both regulated by the hypothalamus. In a zebrafish model reported by Blanchet et al, knock down of the *Myt1l* orthologs (*Myt1la* and *Myt1lb*) resulted in loss of oxytocin (OXT) expression in the neuroendocrine preoptic area of the hypothalamus; moreover, arginine vasopressin (AVP) expression was also significantly decreased in the neuroendocrine preoptic area but not in the ventral hypothalamus (Blanchet et al., 2017). Thus, there is evidence for hypothalamic dysregulation in organisms with *Myt1l* mutations, which raises the possibility that MYT1L haploinsufficiency might result in hypothalamic dysfunction in the mammalian brain as well.

The hypothalamus is a key brain region involved in the regulation of social behaviors, including maternal care and aggression, as well as motivated behaviors such as food intake that can lead to weight gain; behaviors that are frequently disrupted in neurodevelopmental conditions such as autism and Rett syndrome (Shimogori et al., 2010). Oxytocin-producing neurons of the paraventricular nucleus of the hypothalamus (PVN) have long-range projections to multiple brain areas as well as dendrites that release neuropeptides locally (Francis et al., 2014; Neumann, 2007; Seoyoung Son et al., 2022). A very similar peptide, AVP is also synthesized in the same nuclei, plus parts of the limbic system in males (Francis et al., 2014). Both neuropeptides are known to be involved in social behaviors such as social attachment, social recognition, maternal behavior, communication and aggression (Hammock, 2015). Animal studies have suggested that OXT exposure is related to improved social skills and lower reactivity to stressors. OXT dysregulation has been suggested to play a role in multiple neurodevelopmental conditions, such as autism, Williams syndrome, Fragile X syndrome, and Prader Willi syndrome, all of which present with significant social difficulties (Francis et al., 2014; Keech et al., 2018). The role of the hypothalamus has been studied in other genetic mouse models of liability for monogenic neurodevelopmental disorders. In a mouse model of Rett syndrome, targeted loss of MECP2 in *Sim1*-expressing neurons (paraventricular, supraoptic, and posterior hypothalamic nuclei, lateral olfactory tract of the amygdala) resulted in an abnormal physiological stress response, aggression, hyperphagia, and obesity. Of note, expression of AVP and OXT was normal in these mice, which suggests that the noted behavioral effects of hypothalamic dysfunction might not always be mediated by deficiency of these neuropeptides (Fyffe et al., 2008). Likewise, in a mouse model for Smith-Magenis syndrome, conditional knockout of *Rai1* in *Sim1-*positive neurons resulted in obesity due to hyperphagia, a feature frequently seen in affected humans (Huang et al., 2016).

The above supports a key role for neurodevelopmental syndrome genes in the hypothalamus for certain sets of behavior. However, before conducting such conditional genetic studies in MYT1L, we first sought to understand the hypothalamic and hypothalamus-mediated phenotypes in a global heterozygous *Myt1l* mutant (Het) animal. We aimed to determine if *Myt1l* heterozygosity had similar molecular consequences in the hypothalamus as reported for other regions (Chen et al., 2021; Kim et al., 2022; Wöhr et al., 2022; Yen et al., 2024), and if OXT and AVP cell counts were perturbed as had been shown in zebrafish (Blanchet et al., 2017). We then examined a variety of behavioral and metabolic phenotypes, with a particular focus on those previously linked to OXT, to determine if they were also disrupted. If so, this would suggest that MYT1L haploinsufficiency in the hypothalamus may be related to the obesity, hyperphagia, and social behavior abnormalities frequently seen in patients (Blanchet et al., 2017; Coursimault et al., 2022). We found that the *Myt1l* Het hypothalamus did show a similar scale of molecular disruption to previously studied brain regions, and some corresponding alterations in OXT and AVP cell count, though not completely replicating the direction of effect of the observations in zebrafish. However, most OXT-associated behaviors studied here were unchanged, precluding any OXT-focused rescue studies. We did observe an interaction between diet and genotype on weight gain. Finally, there was an absence of major metabolic, pubertal, and other signatures of gross endocrine dysfunction, suggesting that these autonomic systems are largely intact in the context of MYT1L haploinsufficiency.

## Methods and Materials

### Animals

All experimental protocols were approved by and performed in accordance with the relevant guidelines and regulations of the Institutional Animal Care and Use Committee of Washington University in St. Louis and were in compliance with US National Research Council’s Guide for the Care and Use of Laboratory Animals, the US Public Health Service’s Policy on Humane Care and Use of Laboratory Animals, and Guide for the Care and Use of Laboratory Animals.

All mice used in this study were maintained and bred in the vivarium at Washington University in St. Louis. Generation of the MYT1L haploinsufficiency model was described previously (Chen et al., 2021). Breeding pairs for experimental cohorts comprised *Myt1l* heterozygous mutants (Hets, JAX Stock No. 036428) and wild type (WT) C57BL/6J mice (JAX Stock No. 000664) to generate male and female *Myt1l* Het and WT littermates. For all experiments, adequate measures were taken to minimize any pain or discomfort. The colony room lighting was on a 12:12h light/dark cycle (lights on 6am to 6pm); room temperature (∼20-22°C) and relative humidity (50%) controlled automatically. Mice were housed in static (28.5 x 17.5 x 12 cm) translucent plastic cages with corncob bedding, Nestle squares and a cardboard tube (Bio-Tunnels, Bio-Serv). Standard lab diet and water were freely available. Upon weaning at postnatal day (P)21, mice for behavioral testing were group-housed according to sex and genotype.

### ATAC-seq

ATAC-seq was performed as previously described (Chen et al., 2021). Hypothalamus for 5 (3F, 2M) WT and 5 (3F, 2M) Het P60-70 mice were dissected and homogenized in cold nuclear isolation buffer. Lysates were filtered through a 40 μm mesh strainer and spun down. 100,000 nuclei were added to the tagmentation reaction (Illumina Tagment DNA TDE1 Enzyme and Buffer Kit, 30 minute incubation at 37°C) for each sample. DNA fragments were purified and ligated with sample-specific adaptors (QIAquick PCR Purification Kit, Phusion primers). Libraries were purified (AMpure beads at 1:1.8 dilution), assessed for quality, and submitted to GTAC Washington University School of Medicine for Novaseq (50M reads per sample).

### DAR and motif enrichment analysis

Low peak counts were filtered out using the edgeR package and excess variation was removed via the RUVseq package. We then normalized the peak counts for DNA composition and generated a log2(cpm) table storing the count data. The data was fitted into a likelihood ratio test to identify differentially accessible regions across genotypes using edgeR. Significant DARs were determined by FDR < 0.1 cutoff (**Supplemental Table 1**). The peaks were categorized into TSS and non-TSS, with peaks ± 1kb away from a gene transcription start site defined as TSS and the rest as non-TSS. The non-TSS peaks were associated with genes using GREAT via the single nearest gene approach (5kb) and curated regulatory domains included (McLean et al., 2010).

### Motif analysis

Motif analysis was performed on significant DARs (FDR < 0.1) separately for TSS and non-TSS peaks, using the R package monaLisa (v1.0.0) (Machlab et al., 2022). For known motif enrichment analysis, the JASPAR2020 motif database was used for motif finding in both less and more accessible regions, with all ATAC-seq peak regions as background (Rauluseviciute et al., 2024). For de novo motif discovery, the k-mer unbiased motif analysis was performed to search for the MYT1L core binding motif using 5 as the k-mer length and all ATAC-seq peaks as background. Selected significantly enriched motifs (FDR < 0.0001) were visualized through the monaLisa package function plotMotifHeatmaps.

### RNA-seq

RNA-seq was performed as previously described (Chen et al., 2021). Hypothalamus for 6 (3F, 3M) WT and 6 (3F, 3M) Het P60-70 mice were dissected and RNA extracted. Total RNA integrity (RIN) was determined with Agilent 4200 Tapestation. Libraries were prepared using 10ng total RNA with RIN > 8.0. ds-cDNA was prepared according to manufacturer protocol (SMARTer Ultra Low RNA kit for Illumina Sequencing, Takara-Clontech). cDNA was fragmented, blunt-ended, and A base, adaptor-ligated. Fragments were amplified for 15 cycles with unique dual index tags and sequenced using paired end reads extending 150 bases (Illumina NovaSeq-6000).

Sequencing reads were trimmed (Trimmomatic, (Bolger et al., 2014)) and assessed for quality (fastQC, (Andrews, 2024)). Reads were filtered (Bowtie2, (Langmead and Salzberg, 2012)) and mapped (STAR, (Dobin et al., 2013)) to the mouse genome (mm10, (O’Leary et al., 2024)). Quantification was performed on individual samples using HTSeq (Putri et al., 2022).

### DGE analysis

Differential RNA-seq analysis was performed for RNA dissected from the hypothalamus of P60-70 WT control and MYT1L heterozygous mice. We first checked the library size, read count distribution, BCV value, Pearson correlation, hierarchical clustering, multidimensional scale plots to make sure there were no obvious outlier samples. Then, the raw read counts were normalized for RNA composition and unwanted variation was removed using the R packages edgeR (v3.36.0) and RUVseq (Risso et al., 2014; Robinson et al., 2010). Differential gene expression analysis was performed using edgeR and a likelihood ratio test was applied to identify differentially expressed genes (DEGs) across genotypes. P-values were adjusted to correct for multiple testing using the Benchamini-Hochberg false discovery rate (FDR) method. DEGs with FDR < 0.1 were considered to be significant and subjected to downstream analyses (**Supplemental Table 1**). Heatmap and volcano plot for DEGs were generated by R function heatmap.2 from the gplots package and R package EnhancedVolcano respectively (Blighe, 2024; Galili, 2024).

### Gene ontology analysis

Differentially expressed genes identified through the aforementioned DGE analysis were used to assess significantly enriched biological processes and pathways using Cytoscape BiNGO analysis tool (Maere et al., 2005). The hypothalamus-specific genes identified via P60-70 hypothalamus RNA-seq were used as the reference gene set. *p* values were adjusted using Benjamini-Hochberg FDR correction and FDR < 0.05 cut-off was used to define significantly enriched pathways.

### Gene set enrichment analysis

Gene set enrichment analysis (GSEA) was performed using GSEA (v4.3.2) (Subramanian et al., 2005). We first examined a collection of ontology gene sets available on the Molecular Signatures Database (MSigDB) to obtain a comprehensive overview of the biological pathways impacted by the loss of MYT1L. Then, we used the Notch signaling pathway, negative regulation of Wnt signaling pathway, and neuropeptide hormone activity gene sets downloaded from MSIgDB and the pre-defined ‘early-fetal’ and ‘mid-fetal’ gene sets as input for our final GSEA (**Supplemental Table 1**) (Kang et al., 2011; Katayama et al., 2016). All analyses were performed with ‘‘gene_set’’ as permutation type and 1,000 permutations. The gene sets with an FDR < 0.1 were considered significantly enriched.

### Integration of RNA-seq and ATAC-seq data

The DEGs and DAR-associated genes, both TSS and non-TSS, were overlapped together to identify genes with both significantly altered gene expression and chromatin accessibility (**Supplemental Table 1**). The direction of regulation was taken into consideration as well. The overlap was determined to be significant with Fisher’s Exact Test, p-value<0.05.

### Data Availability

ATAC-seq and RNA-seq have been deposited to Gene Expression Omnibus (GEO) at accession number GSE173589.

### Immunofluorescence

To characterize cell numbers in the PVN expressing AVP and OXT, we employed immunofluorescence techniques as previously described(Mulvey et al., 2023). Specifically, brains of PND 80-100 mice (Het n=6 males, 6 females; WT n= 6 males, 5 females) were removed following isoflurane overdose and fixed in paraformaldehyde in PBS (4%) followed by serial sucrose in PBS solutions (15%, 30%). Following post-fixation, brains were embedded in OCT (Sakura, Torrence CA) and sectioned at 40 µm using a Leica CM1950 cryostat (Buffalo Grove, IL) and processed for immunofluorescence as free-floating sections. For immunofluorescence, sections were incubated in 1x PBS with 5% normal donkey serum and 1:500 rabbit anti-OXT (1:500, Millipore, AB911) and 1:500 mouse anti-AVP (1:500, Santa Cruz Biotech, sc-9888). Slices incubated in primary antibodies overnight at 4C on a horizontal mixer, rinsed three times in 3xPBS, followed by 1 hour of incubation in 1X PBS with 5% normal donkey serum and 1:1000 each of Alexa Fluor 488 donkey-anti-rabbit, and Alexa Fluor 568 donkey-anti-mouse. Sections were rinsed again with 1xPBS, then incubated for 5 minutes with 1:20,000 DAPI for nuclear fluorescence, rinsed with 1xPBS and then mounted on slides. Slides were covered by nail polishing coverslips after an application of Prolong Gold. Slides were stored in light-proof slide boxes at 4C until Z-stack imaged on a Leica microscope (Thunder DMi8) at 20x resolution.

Image editing was performed using Fiji by ImageJ software and only included re-scaling of resolution, brightness and contrast. Fluorescent Z-stack images of DAPI were stacked and used to create a composite of the nucleated PVN of the hypothalamus to measure the total area of the PVN in mm^2 for normalization measurements of cell counts. Fluorescent images of AVP and OXT were merged using ImageJ. Following merging, cell counts of AVP and OXT cells were measured. This cell count was normalized to the PVN area to result in %AVP or %OXT cells /mm^2.

### Adrenalectomies and Gonadectomies

As a gross measure of endocrine system integrity, we assessed adrenal and gonadal weights in Het and WT littermates based on previously published methods (Finco and Hammer, 2018; Idris, 2012; Mason et al., 2018). To do this, we anesthetized 55 WT (25M, 30F) and 51 *Myt1l* Het (27M, 24F) littermates using 1.5-2% isoflurane. Ages (days) at dissection were the following (Median [Range]): WT male (82 [59-110]), WT Female (87, [62-110], Het male (84, [59-110]), Het female (90, [62-110]). Mice were sacrificed by decapitation. Females were dissected for both ovaries and adrenal glands, while males were dissected for both testes and adrenal glands. Dissected tissue was left to dry at 4C for 48 hours. Organs were weighed on a high-sensitivity digital scale (min 1mg, max 120g, d = 0.1mg). Left and right adrenal glands were weighed separately and combined for graphing. Left and right gonads were weighed together. Body weights were also taken immediately prior to anesthesia administration.

### Behavioral Tasks

Hypothalamically-mediated behavioral function was assessed in five independent cohorts of mice (**Table 1**). Mice were handled for 3-5 days prior to the behavioral task and the tails of mice were marked with a non-toxic, permanent marker during weight collection and regularly thereafter to easily distinguish mice during testing. For all tasks, the mice were acclimated to the testing room at least 30 minutes prior to the start of testing. Testing orders were randomly counterbalanced for genotype across apparatuses and trials. All assays were conducted by female experimenters blinded to experimental group designations during testing, and all testing occurred during the light/inactive phase.

**Table 1.** Order of and age at behavioral testing per cohort with sample size.

| Cohort | Behavioral assessment | Age | Sample size |  |  |
| --- | --- | --- | --- | --- | --- |
|  |  |  | Total | Females | Males |
| 1 | Maternal care behavior | $M=P105$ (range P65-124) | 22 | 22 | -- |
| 2 | Social dominance | P110 | 48 | 24 | 24 |
|  | Resident intruder | P128 | 24 | -- | 24 |
| 3 | Sexual Maturity | P14-P50 | 54 | 29 | 35 |
| 4 | Metabolic assessments | P25 & P120 | 75 | 33 | 32 |
|  | Feeding behavior | P28-P35 & P123-P130 | 36 | 18 | 16 |
| 5 | Metabolic assessments | P120 | 32 | 12 | 20 |
|  | Feeding behavior | P123-P130 |  |  |  |

#### Maternal Care Behavior

The impact of MTY1L mutation on maternal care was tested in 22 dams. WT (n=12) and MYT1L (n=10) females were bred to C57BL/6J WT males and maternal care was recorded with their first litter at P2 and P4. At time of assessment, the WT dams aged P91.5 (on average, range P65-105) and the Het dams aged P114.5 (on average, range P70-134). The full breeding cage was brought into the testing room 30 minutes prior to testing. The dam and sire were removed from the cage and placed in a clean cage for 10 minutes, while the cage with pups that maintained the nest was placed on a heating pad. After 10 minutes, the pups were evenly placed in the three corners of the cage outside of the nest and the dam was immediately returned to the cage and filmed (Sony HDR-CX560V camera) for 10 minutes. If all pups were not retrieved to the nest at the end of 10 minutes, they were returned gently by the experimenter. Fiji(ImageJ) used to determine the area of the nest during the first minute of pup retrieval and again during the last minute of recording. Ethovision 14XT (Noldus) was used to track the movements and behaviors of the dam, including dam self-grooming and pup grooming, entries, exits and time in the nest. Pup retrieval latency was manually scored.

#### Social Dominance Tube Test

Between six to eight weeks of age, laboratory mice acquire dominance behaviors to create social hierarchy ranks within their cage environments, or social groups. We used the social dominance tube test (**Figure 5A**) to assess social dominance ranks within cages at P110 to identify any influence of MYT1L haploinsufficiency on cage rank dynamics, adapted from our previously published methods (Chen et al., 2021). All mice were housed in groups of three. Briefly, each mouse was habituated to the testing tube on days 1 and 2 by allowing it to move completely through the tube starting from one side on each day. On each of the three test days, each mouse in a cage was paired in the tube test with each of its cage mates to determine hierarchy rank from most dominant (ranked 1, dominated the other two cage mates) to most submissive (ranked 3, submitted to the other two cages mates, **Figure 5B**). The dominance bouts were video recorded and subsequently scored for a submissive mouse (first to back out with all four paws leaving the tube) and a dominant mouse (held its ground or pushed the other mouse out of the tube).

#### Resident Intruder

To evaluate aggressive and agonistic behaviors, we assessed male mice in the resident intruder paradigm at P128 following our previously published methods (Nygaard et al., 2023). To establish a territory, male mice were isolation-housed and the cages remained unchanged for ten days prior to testing. A sexually naive strain- and age-matched female was added to each male cage on day 6 and removed 24 hours prior to testing to further potentiate territorial behaviors. Intruder mice were strain- and age-matched novel male mice bred in the same colony space. For testing, an intruder mouse was added directly into the resident home cage in a sound-attenuating box under white light and allowed to freely interact for 10 minutes. Interactions were filmed with a digital camera (Sony HandyCam). At the end of testing, the intruder was removed and placed in a clean holding cage until all cage-mates complete testing. This was repeated over two additional days with a novel intruder mouse. We monitored interactions for any attacks drawing blood, which would necessitate prematurely ending the assay. However, we did not observe any attacks at this level.

Pose estimation software (DeepLabCut, version 2.2.1) was used to track body parts in space and time from the videos (Mathis et al., 2018; Nath et al., 2019), building upon a previously built model (Nygaard et al., 2023). Specifically, we labeled 50 frames taken from 72 videos plus an additional 1700 frames of closely interacting animals, and 95% were used for training. We used a dlcrnet_ms5-based neural network with default parameters for 200,000 training iterations. We validated with two shuffles, and found the test error was 7.95 pixels, with the train error at 5.58 pixels (image size was 1920 by 1080). We then used a p-cutoff of 0.6 to condition the X,Y coordinates for future analysis. This network was then used to analyze all videos in the experiment.

The random forest classifier Simple Behavior Analysis (SimBA)(Nilsson et al., 2020) was then used to extract attack behavior from the pose estimation data, building upon a previously built model(Nygaard et al., 2023). Training videos were annotated for attack behavior using the SimBA event-logger using 180 annotated behavior files, downloaded from https://osf.io/tmu6y/ in addition to four in-house annotated videos. All training files were annotated according to definitions found in the simBA preprint (Nilsson et al., 2020; Nygaard et al., 2023). Random forest classifiers were trained using default hyperparameters, and classifier performances were evaluated. We set the discrimination threshold for scramble attacks to 0.55. The minimum bout length was 200 ms. Once classifiers were validated, predictive classifiers were applied to all videos in the dataset. Statistics regarding outcome variables of total time, the number of frames, total number of instances, mean instance interval, and mean interval between behavior instances were generated.

We employed a custom Python script that leveraged X, Y coordinates derived from DeepLabCut pose-estimation for both the nose and tailbase of mice to extract sniffing behaviors exhibited by the animals. A sniff event was operationally defined as the nose point of one mouse being within a spatial proximity of 2 cm or 3.29 pixels to the nose or tailbase points of another mouse.

Distinct categories of sniffing behaviors were identified based on the spatial relationships between the nose and tailbase points of interacting animals. Specifically, the interaction labeled as "Resident Nose to Intruder Nose" was indicative of facial sniffing. Moreover, the interaction denoted as "Resident Nose to Intruder Tailbase" represented instances where the resident mouse initiated anogenital sniffing. Conversely, "Intruder Nose to Resident Tailbase" characterized situations where the intruder mouse initiated anogenital sniffing. These categorizations facilitated a comprehensive analysis of the various sniffing patterns observed during social interactions of the resident intruder paradigm

#### Sexual Maturity

Onset of sexual maturity was checked every other day starting at P14, based on the literature (Caligioni, 2009; Hoffmann, 2018). This was determined by prepubital separation in males and vaginal opening in females. On each assessment day, In females, a sterile cotton swab was used to gently check for vaginal opening. The male mice were held by the tail and the skin surrounding the penis was gently pushed back. The prepubital region was marked as either open or closed. Each mouse was then weighed and returned to its home cage. Determination of pubertal onset and weight continued for 4 days after onset.

#### Feeding Behavior (FED3)

The impact of *Myt1l* mutation on feeding behavior was assessed in multiple cohorts (**Table 1**; **Figures 7A & 10F**) across seven days. To chow pellet achievement across time, mice were singly housed and a Feeding Electronic Devices (FED3) (Matikainen-Ankney et al., 2021; Nguyen et al., 2016) was placed in the home cage. The FED3 is a wireless, battery-powered device designed for home cage training of mice in operant tasks. Mice interacted with the FED3 through two nose-poke holes, and the FED3 responded to the mice with visual and auditory stimuli, and food pellets (grain pellets, Bio-SERV) are dispensed into a trough. Behavior was assessed in four phases: magazine training (24 hours), dual active (24 hours), fixed ratio 1 (FR1; 72 hours), and resetting progressive ratio (48 hours; **Figure 8A**). During magazine training, the device continuously released a pellet once the previous one was removed from the trough. No hole poking was required. Release of a new pellet was accompanied by lights and a tone. During dual active training, both the left and right hole were active and a poke in either resulted in light and sound signal and release of a pellet. Nose poke holes were inactive when a pellet is already present in the trough. During FR1, the right hole was inactive and only a poke into the left hole resulted in the lights, tone and release of a pellet. A poke into the inactive hole did not result in any signal from the device. During the final stage, resetting progressive ratio 1 (PR), the number of nose pokes into the active hole increased by one with each subsequent pellet. After 30 minutes of inactivity, the counter reset to one, with one nose pokes releasing one pellet. Each pellet removed from the device is quantified as “achieved” and not “consumed” because, while we think the animals are eating the pellets and did not observe pellets remaining in the cage, we cannot know for certain. The pellets could potentially be ground down with the nesting and other materials in the cage. Animals were removed from testing if they did not interact with the device in the first 24 hours of testing. Devices were cleaned and reset in between animals. PR data for two mice were lost due to equipment failure during testing.

#### High Fat Diet Exposure

Starting at weaning, a subset of mice were fed either regular chow (10% fat, 20% protein, 70% carbohydrate, 3.10kcal/g) or a moderately high fat diet (HFD; 45% fat, 20% protein, 35% carbohydrate, 4.73kcal/g; D12451 formula Research Diets, Inc) for 14 weeks until P120. Mice were sex- and genotype-match housed 2-4 mice per cage. Food was provided in custom hanging food receptacles, measuring 8.25cm in diameter and 5.08 cm deep with a round metal mesh cap to ensure large pellets could not be removed but not impede feeding. Each receptacle was filled with 100g of food twice per week, and weighed before and after. Average consumption per mouse was determined by calculating the difference in food weight divided by the number of days in the cage and then by the number of mice per cage. Mice were also weighed at time of food change.

### Biochemical Analysis of Plasma

At both P25 and P120 (**Figure 7A**), plasma samples were acquired to measure levels of glucose, cholesterol, triglycerides, and free fatty acids through the Animal Model Research Core in the Nutrition Obesity Research Center at Washington University School of Medicine in mice that were 12 hr food restricted and those that were not. Approximately 40-50 uL of blood was collected by tail vein bleed with a heparinized capillary tube (Fisher, 22-260-950) which was then spun down at 6000 rpm for 15 min at 4C. The 15 uL of plasma supernatant was then extracted and stored at -80C until processing. Total triglycerides, total cholesterol, glucose, and free fatty acids in plasma were measured using the following kits: Triglyceride Infinity Kit (Thermo Scientific, catalog no. TR22421), Cholesterol Infinity Kit (Thermo Scientific, catalog no. TR13421), Autokit c, glucose diagnostic reagent (Fujifilm Wako, catalog no. 99703001) and NEFA (Fujifilm Wako, catalog no. 99934691). For each kit, 1 ul of plasma sample or standard was added to 100 ul reagent, mixed and the reaction incubated for 30 mins at room temp. Total triglycerides and glucose were determined by readings at A540, and total cholesterol was read at A490. Free fatty acids in plasma were determined in a two-step reaction. First, 2 volumes (50 ul) of reagent A were aliquoted to wells, then 1 ul of plasma or standard added and incubated for 15 min at room temperature. Then, color was developed with the addition of 1 volume (25 ul) of reagent B to all wells. The plate with samples and standards was then incubated an additional 15 minutes at room temperature and read at 540 nm. The sample concentrations were calculated from a standard curve correcting for blanks and a secondary wavelength at 660 nm.

### Statistical Analysis

Behavioral statistical analyses and data visualization for data were conducted using IBM SPSS Statistics (v.28 and v.29). Prior to analyses, data were screened for missing values and fit of distributions with assumptions underlying univariate analysis. This included the Shapiro-Wilk test on z-score-transformed data and qqplot investigations for normality, Levene’s test for homogeneity of variance, and boxplot and z-score (±3.29) investigation for identification of influential outliers. Means and standard errors were computed for each measure. Analysis of variance (ANOVA), including repeated measures, were used to analyze data where appropriate, and simple main effects were used to dissect significant interactions. Sex was included as a biological variable in all analyses. Where appropriate, the Greenhouse-Geisser or Huynh-Feldt adjustment was used to protect against violations of sphericity for repeated measures designs. Multiple pairwise comparisons were subjected to Bonferroni correction, where appropriate. For data that did not fit univariate assumptions, non-parametric tests were used or confirmed parametric results. The Chi-square was used to assess relationships between categorical variables and one-sample t-tests were used to assess differences from chance levels. Sex*genotype effects are reported where significant, otherwise data are discussed and visualized collapsed for sex, with details of all statistical tests and results reported in Table 2. The critical alpha value for all analyses was p < .05 unless otherwise stated. Figure illustrations were generated using BioRender. The behavior datasets generated and analyzed during the current study are available from the corresponding author upon reasonable request.

**Table 2.**
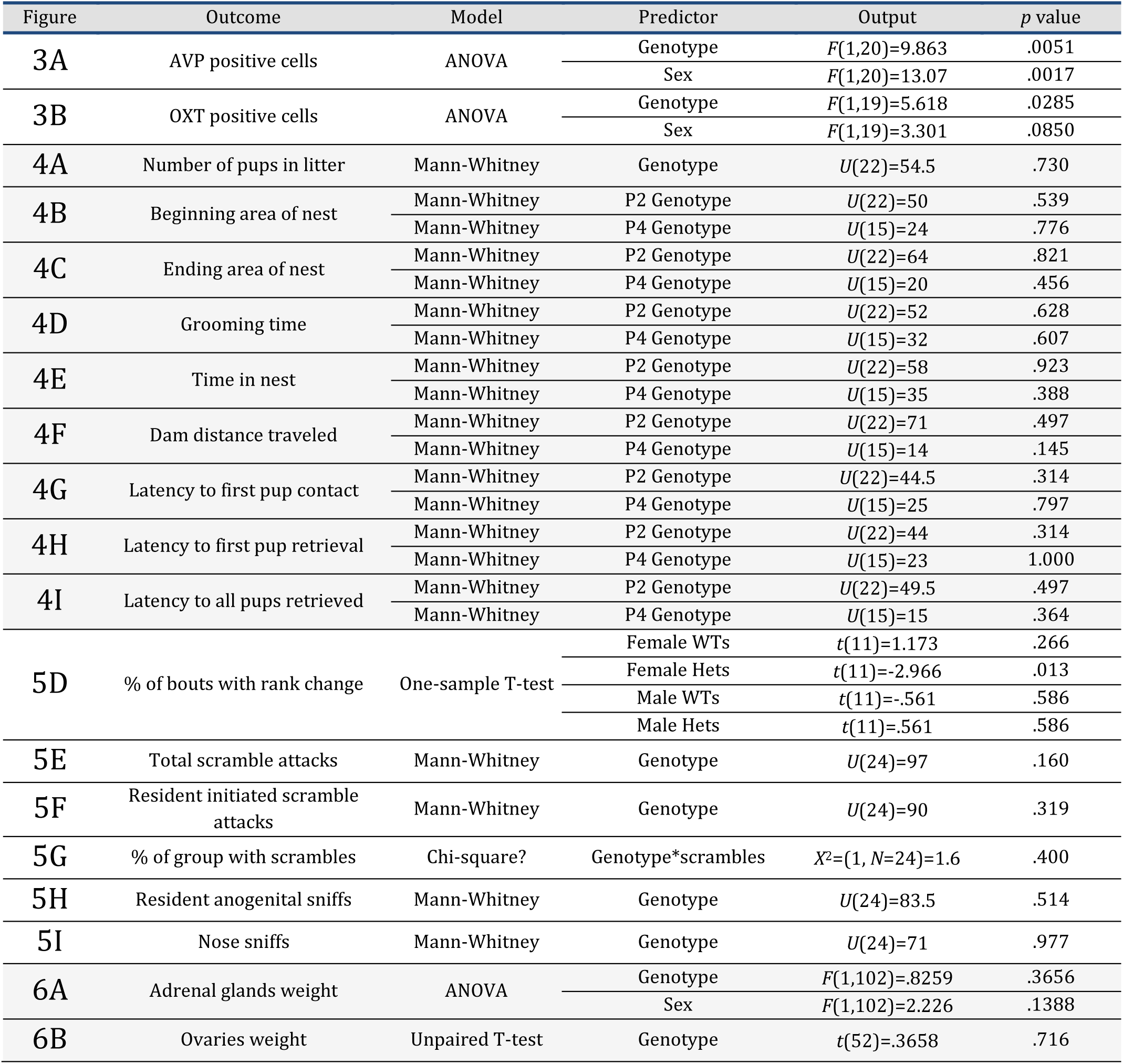
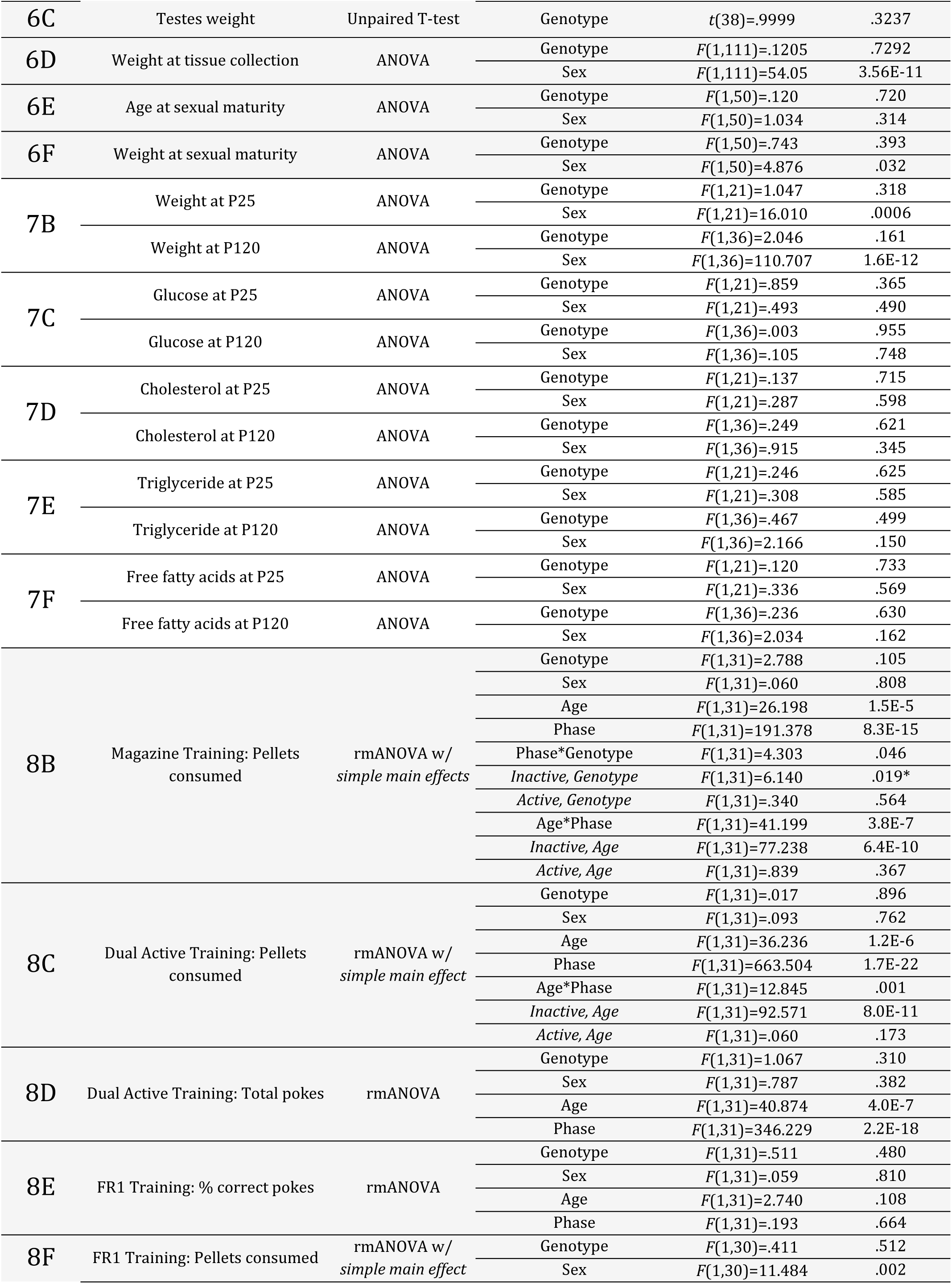
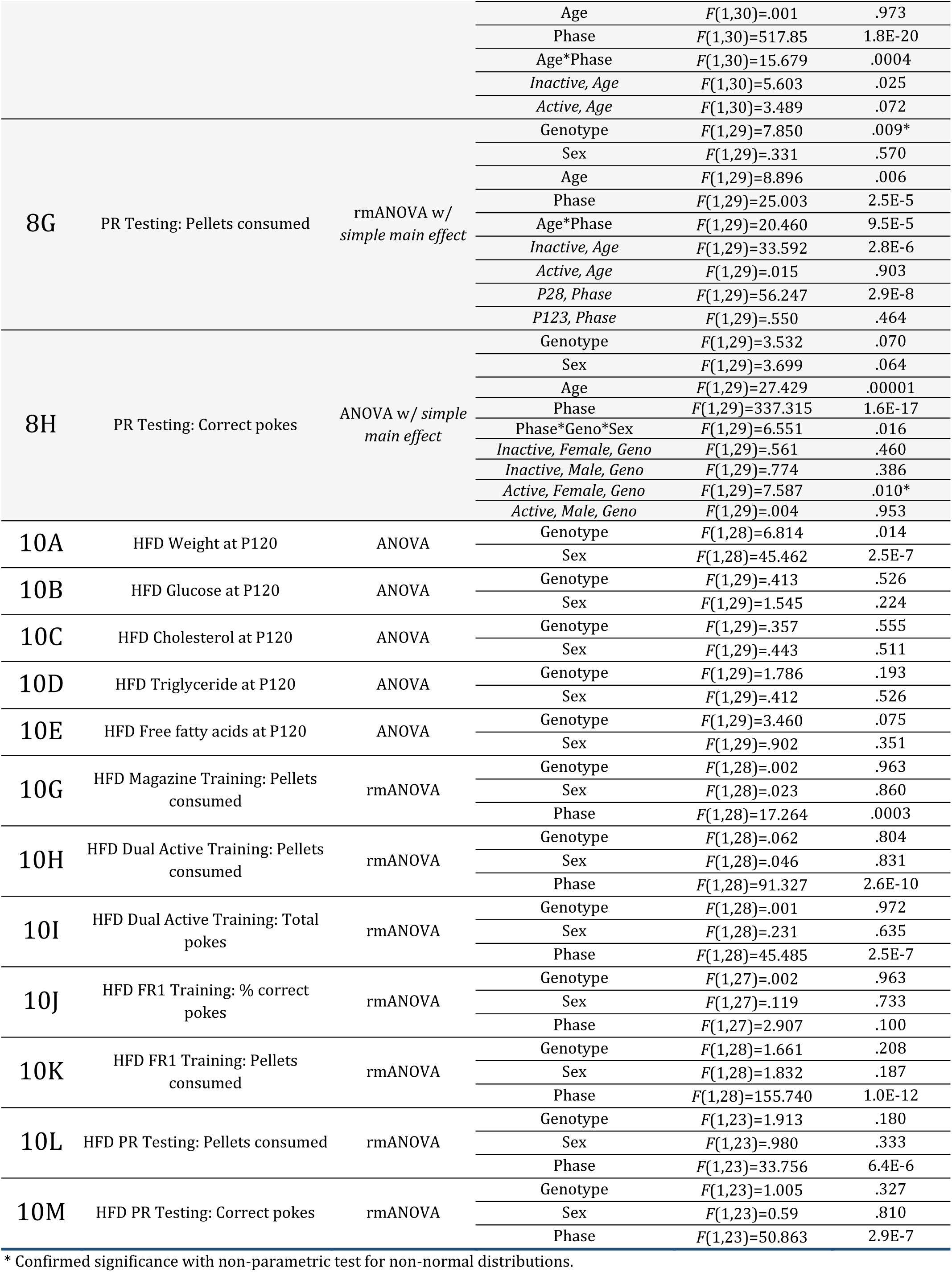
Statistical analysis results.

## Results

### Epigenetic and transcriptomic characterization of MYT1L mutant hypothalamus

Based on MYT1L’s role as a transcription factor (TF), we first set out to determine if MYT1L haploinsufficiency had a clear consequence on chromatin state and gene expression in the hypothalamus with an assessment of open chromatin (ATAC-seq) and gene expression (RNA-seq). We performed ATAC-seq on both Het and WT hypothalami and identified that 12,934 out of 234,478 regions had significantly altered chromatin accessibility **(Figure 1A-C, Supplemental Table 1)**. To determine if these effects were direct or indirect, we examined both known TF binding and novel motif enrichment in the identified differentially accessible regions (DARs). Interestingly, the known MYT1L core binding motif, AAGTT, was not significantly enriched in the differentially accessible promoter or enhancer regions. This suggests that most DARs in the Het hypothalamus are due to indirect actions of MYT1L. Indirect effects of MYT1L in the hypothalamus may include interactions with other cofactors that bind DNA or downstream secondary effects of gene expression. Indeed, there was a significant enrichment of general transcription factor binding motifs in the differentially accessible promoter regions **(Figure 1D)**. Binding motifs of both annotated transcriptional activators (e.g., ELF4 and KLF14) and repressors (e.g., ZBTB7A and THAP11) were enriched. In contrast to the promoter regions, the differentially accessible enhancer regions were more significantly enriched for neurogenic TF motifs (e.g., NEUROD and MEF2 protein families) and activity-dependent TF motifs (e.g., JUND) **(Figure 1E)**. This is similar to prior observations in the mouse cortex, suggesting that MYT1L haploinsufficiency results in altered activity-dependent transcription across the brain (Chen et al., 2021). These findings indicate that MYT1L haploinsufficiency alters chromatin in the hypothalamus and suggests that MYT1L likely acts upstream or in cooperation with other neurogenic co-factors at enhancer regions.

**Figure 1:**
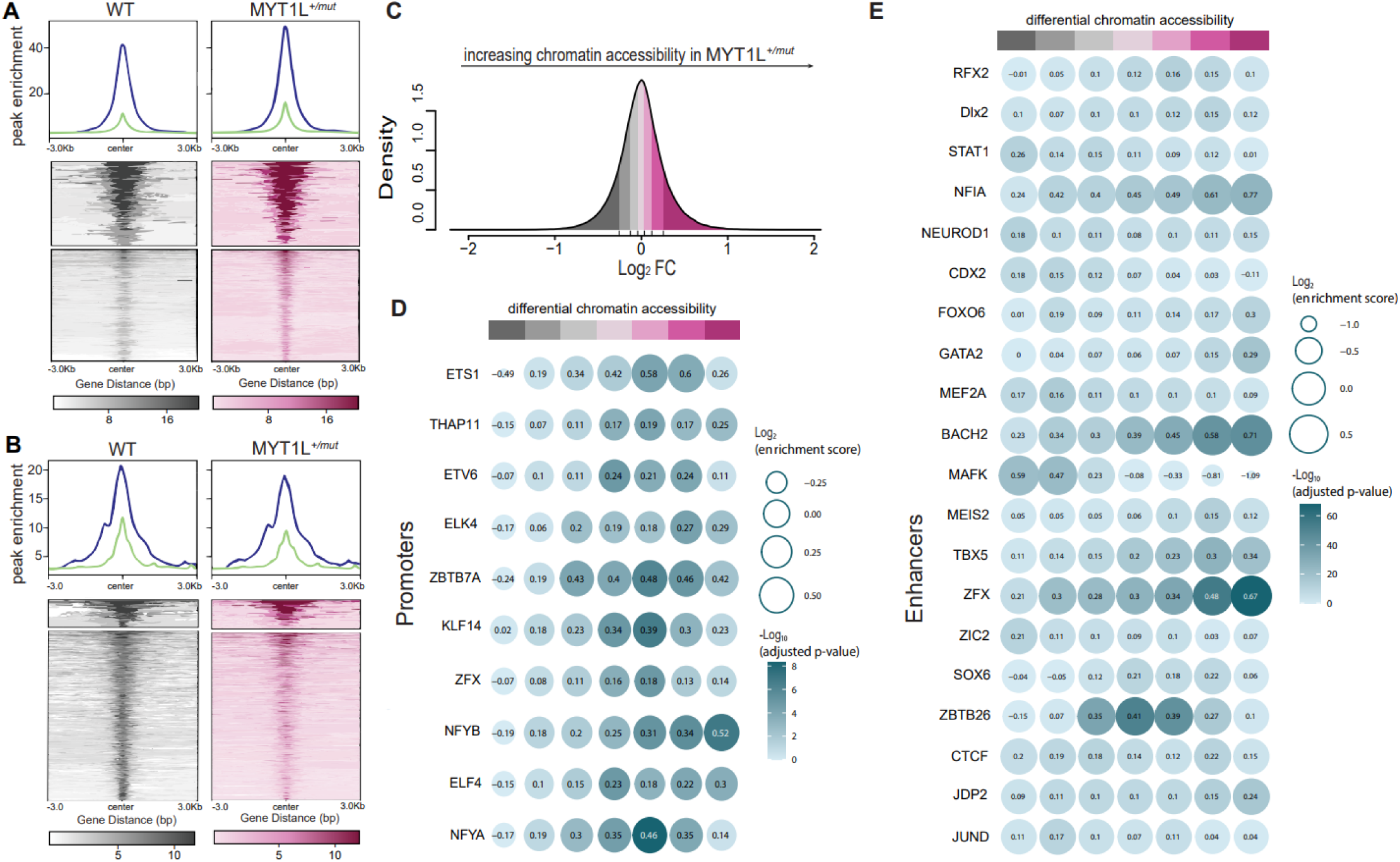
MYT1L mutation altered chromatin accessibility in murine hypothalamus. (A-B) Density plots (top) and genomic heatmaps (bottom) showing differential chromatin accessibility between WT (left) and MYT1L heterozygous (het) mutant mice (right) identified by ATAC-seq (FDR < 0.1) ranging from 3kb upstream of transcriptional start site (TSS) and 3kb downstream of transcriptional end site (TES). (C) Histogram displaying density of chromatin accessibility changes when comparing WT to MYT1L mutant het mice where increasing accessibility in het mice is represented by increasing log_2_FC values. (D-E) Dot plots showing results from monaLisa motif analysis where the size of the dot relates to the log2 enrichment score and the color represents –log_10_ adjusted p-value. Enrichment scores are arranged along the x-axis according to differential chromatin accessibility as in C. In accessible promoter regions (D) we see enrichment of general TF binding motifs while for differentially accessible enhancer regions (E) we see enrichment for neurogenic and activity-dependent TF binding motifs.

Next, we aimed to investigate the downstream transcriptional consequences of MYT1L haploinsufficiency using RNA-seq. From 14,457 genes we identified 718 significantly (FDR < 0.1) differentially expressed genes (DEGs), with more genes being upregulated than downregulated in Hets (Supplemental Table 1). With more genes upregulated upon MYT1L loss, we presume that MYT1L behaves more as a transcriptional repressor than activator in the hypothalamus **(Figure 2A)**. This contrasts with previous findings in the adult mouse cortex where MYT1L loss resulted in roughly equal numbers of up and downregulated genes, yet similar to observations in more recent developing brain single cell data (Chen et al., 2023, 2021; Yen et al., 2024). Thus, we show that MYT1L has context- and region-specific roles.

**Figure 2.**
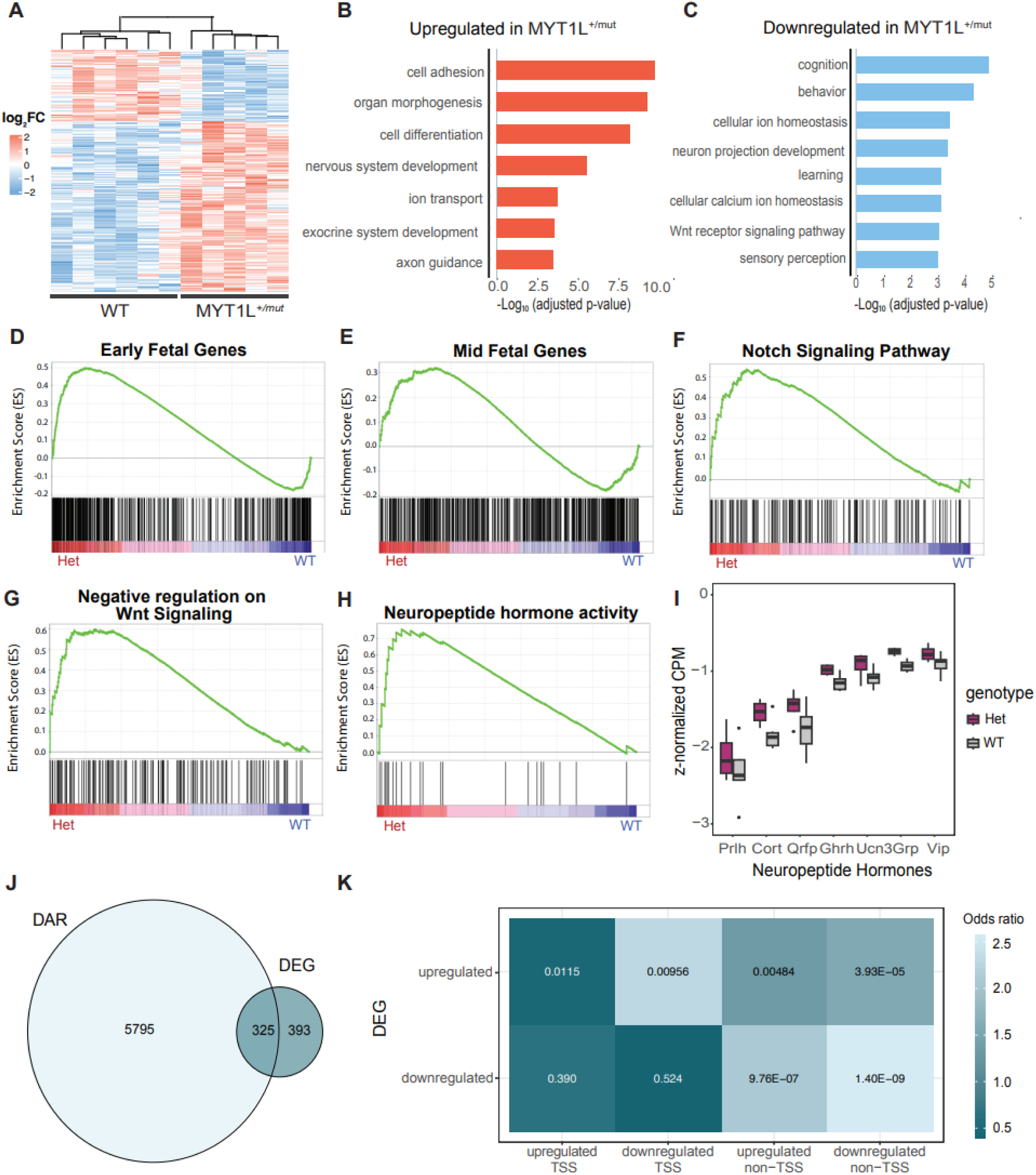
Transcriptional consequences of MYT1L mutation in murine hypothalamus. (A) Heatmap of significant (FDR < 0.1) gene expression changes in Het mice as compared to WT mice with red indicating a higher expression (or Log_2_FC) in Het mice and blue indicating a lower expression in Het mice. (B-C) GO analysis of upregulated (B) and downregulated (C) genes in Hets sorted by – log_10_ adjusted p-value. (D-H) GSEA plots displaying enrichment scores for various categories of genes as they relate to genotype. Green lines indicate the running enrichment score (ES) while x-axis represents the ranked genes in the gene set. (I) Boxplots showing z-normalized CPM values of neuropeptide hormone genes in Het vs WT mice. (J) Venn diagram displaying overlap between differentially accessible chromatin regions from ATAC-seq and differentially expressed genes from RNA-seq. (K) Heatmap showing the correlation between chromatin state alterations (DAR) and gene expression dysregulation (DEG) in promoter (TSS) and enhancer (non-TSS) regions.

To further delineate which biological pathways are activated or repressed upon MYT1L loss, we performed a gene ontology analysis for up- and down-regulated DEGs **(Figure 2B-C)**. With MYT1L loss, general developmental pathways, such as organ morphogenesis and nervous system development, were significantly upregulated **(Figure 2B)**. This is consistent with previous results in the adult cortex (Chen et al., 2021), where the activated developmental pathways are mainly driven by neuronal differentiation and axon guidance genes. This suggests that neurons in Het mice may be retained in a more immature state. In a more direct test of this hypothesis, gene set enrichment analysis (GSEA) was performed and revealed that the Hets had a significant upregulation of early fetal genes, with no change in mid-fetal genes **(Figure 2D-E, Supplemental Table 1)**. Thus, MYT1L may be necessary to suppress early neuronal development programs, echoing MYT1L’s role in prior cortical studies (Chen et al., 2021)). Additionally, genes related to mature functions such as those related to cognition and behavior were downregulated in Hets, driven by reduced expression of genes such as SLC24A2 and EPHA4 **(Figure 2C)**. The loss of MYT1L also dysregulated pathways related to ion transport and homeostasis, whose efficiency is central to synaptic function and neuronal excitability.

Surprisingly, the Wnt receptor signaling pathway, including WNT10A, WNT3 and WNT9B, was significantly downregulated as a consequence of MYT1L loss - in marked contrast to prior studies of MYT1L loss in iPSC derived neurons (Weigel et al., 2023)**(Figure 2F)**. This suggested Wnt downregulation was further supported by the finding that genes involved in ‘negative regulation of Wnt signaling pathway’ were significantly upregulated in Hets on GSEA. Therefore, MYT1L haploinsufficiency may dysregulate Wnt signaling in the hypothalamus. GSEA also showed significant upregulation of Notch signaling pathway genes upon MYT1L loss, consistent with the downregulation of Notch when MYT1L is overexpressed in fibroblasts during transdifferentiation studies (Mall et al., 2017)**(Figure 2G)**.

Due to the crucial role the hypothalamus plays in hormone regulation and secretion, we next examined genes involved in neuropeptide hormone activity using GSEA. Interestingly, MYT1L loss leads to a significant activation of neuropeptide expression, specifically driven by genes encoding or regulating prolactin, cortistatin, and OXT. **(Figure 2H-I)**. Overall, these results suggest that MYT1L loss leads to a dysregulation of neuronal maturation and disrupted neuropeptide systems in the adult mouse hypothalamus.

To evaluate downstream transcriptional effects due to differences in chromatin accessibility caused by MYT1L loss, we integrated both the DARs from ATAC-seq and the DEGs from RNAseq. Out of 718 differentially expressed genes we found 325 that also showed altered chromatin accessibility in response to MYT1L loss **(Figure 2J, Supplemental Table 1)**. To further delineate directional and regional effects of open chromatin on transcription, we compared the overlap of differentially accessible promoters and enhancer regions with up- and down-regulated DEGs. We observed more significant association between chromatin state alterations in enhancers and gene expression dysregulation **(Figure 2K)**. This indicates that MYT1L likely impacts transcriptional processes mostly through altering the accessibility of enhancers, rather than promoters.

### MYT1L haploinsufficiency resulted in altered oxytocin and vasopressin expressing cells in the PVN

Altered neuropeptide RNA levels could correspond to changes in the number of neuropeptide cell numbers, or the amount of RNA produced per cell. Previous literature has identified decreased AVP and OXT expression in a zebrafish model of MYT1L KO in regions analogous to the PVN in rodents (Blanchet et al., 2017). Following our observation that MYT1L loss significantly activates OXT expression, we investigated the impact of MYT1L loss on AVP and OXT expressing cell numbers in the PVN. Furthermore, with sex-specific social impairments in Hets reported previously, we also sought to identify any potential sex effects or sex by genotype interactions. Brain sections were processed for immunofluorescence of AVP/OXT at P80-100 in both WT and Het males and females. AVP/OXT expression was present in the PVN of the hypothalamus of males and females of both genotypes **(Figure 3A,B)**. We found that Het mice had decreased AVP yet increased OXT cell counts in the PVN compared to WT littermates **(Figure 3C,D)**. Furthermore, we identified that female mice had overall decreased AVP cell counts compared to male littermates yet there was no significant sex effect on OXT cell counts **(Figure 3E,F)**. Finally, no significant interaction of sex with genotype was identified for either AVP or OXT counts. Overall, our findings suggest the increased OXT RNA levels may have been driven by a change in OXT cell numbers.

**Figure 3.**
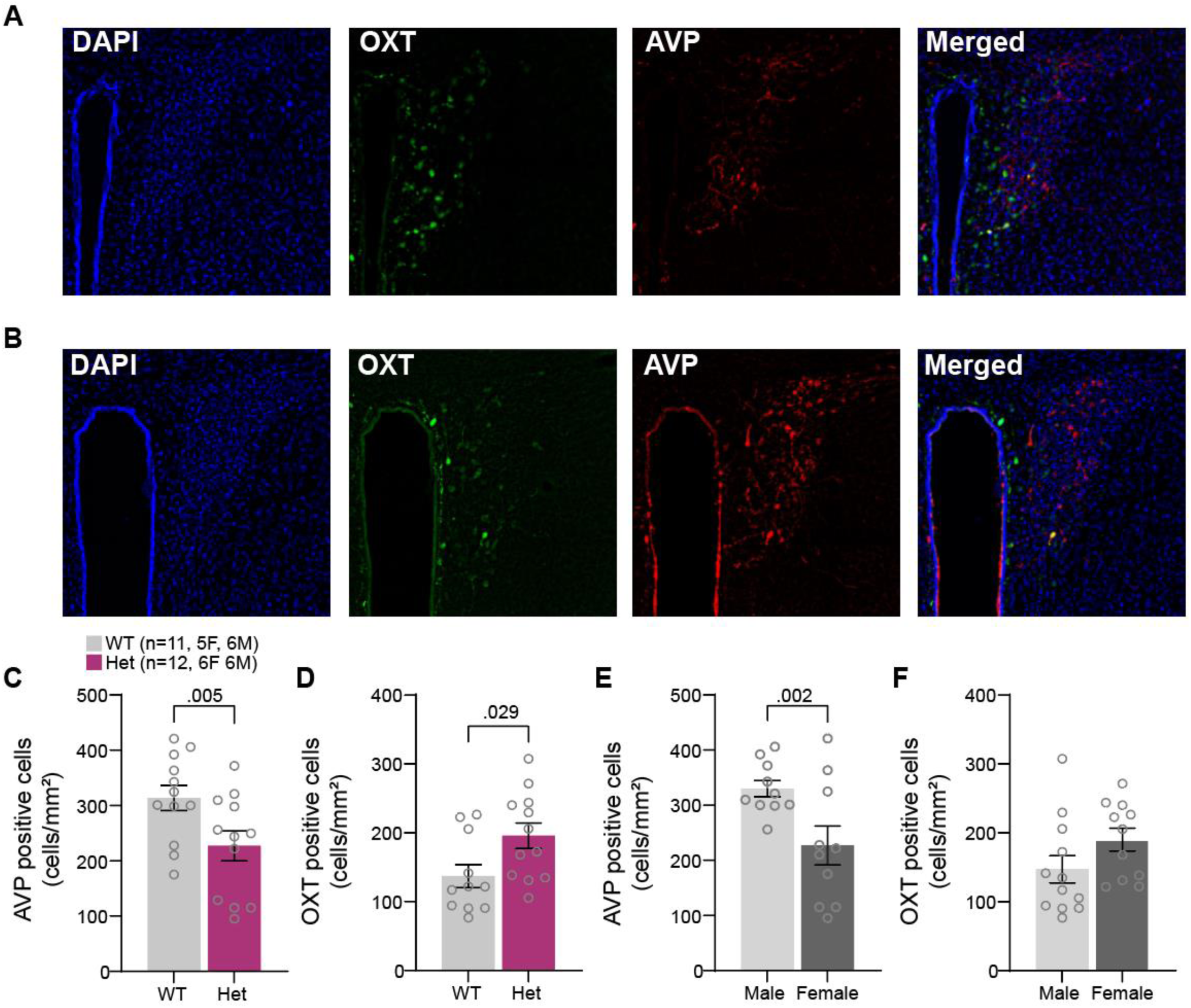
*Myt1l* loss disrupted the number of AVP+ and OXT+ cell numbers in the paraventricular nucleus. (A-B) Immunofluorescence for nuclei (DAPI; blue), oxytocin (OXT; green), arginine vasopressin (AVP; red) in the PVN of the hypothalamus from Het mice (A) and WT mice (B). (C-D) Het mice had significantly less AVP positive cells (C) yet significantly more OXT positive cells (D) compared to WT. (E-F) Females had significantly less AVP positive cells compared to males (E) with no significant sex bias in OXT expressing cells (F). Sample sizes are presented in panel C. Data are represented as mean ± SEM with individual points presented as open circles.

### Female MYT1L haploinsufficiency did not influence maternal behaviors

The neuropeptides OXT and prolactin are critically involved in maternal physiological and behavioral (bonding) responses to newly born offspring (Champagne et al., 2001; Insel et al., 1997; Pedersen et al., 1982; Pedersen and Prange, 1979). To examine the potential impact of upregulation of neuropeptide expression with MYT1L loss, we examined a variety of maternal behaviors in WT and Het dams on postnatal days 2 and 4 of their first litter, including a pup retrieval assay. MYT1L loss in the dam did not impact the number of pups per litter (**Figure 4A**) nor maternal behaviors. Specifically, Het dams performed similarly to WT dams on size of nest, time spent grooming itself or pups, time in nest, and distance traveled around the cage (**Figure 4B-F**). During retrieval of pups, no differences were observed for latency to first contact with a pup, first pup retrieval or last pup retrieval (**Figure 4G-I**). Thus, MTY1L haploinsufficiency did not impact maternal behaviors.

**Figure 4.**
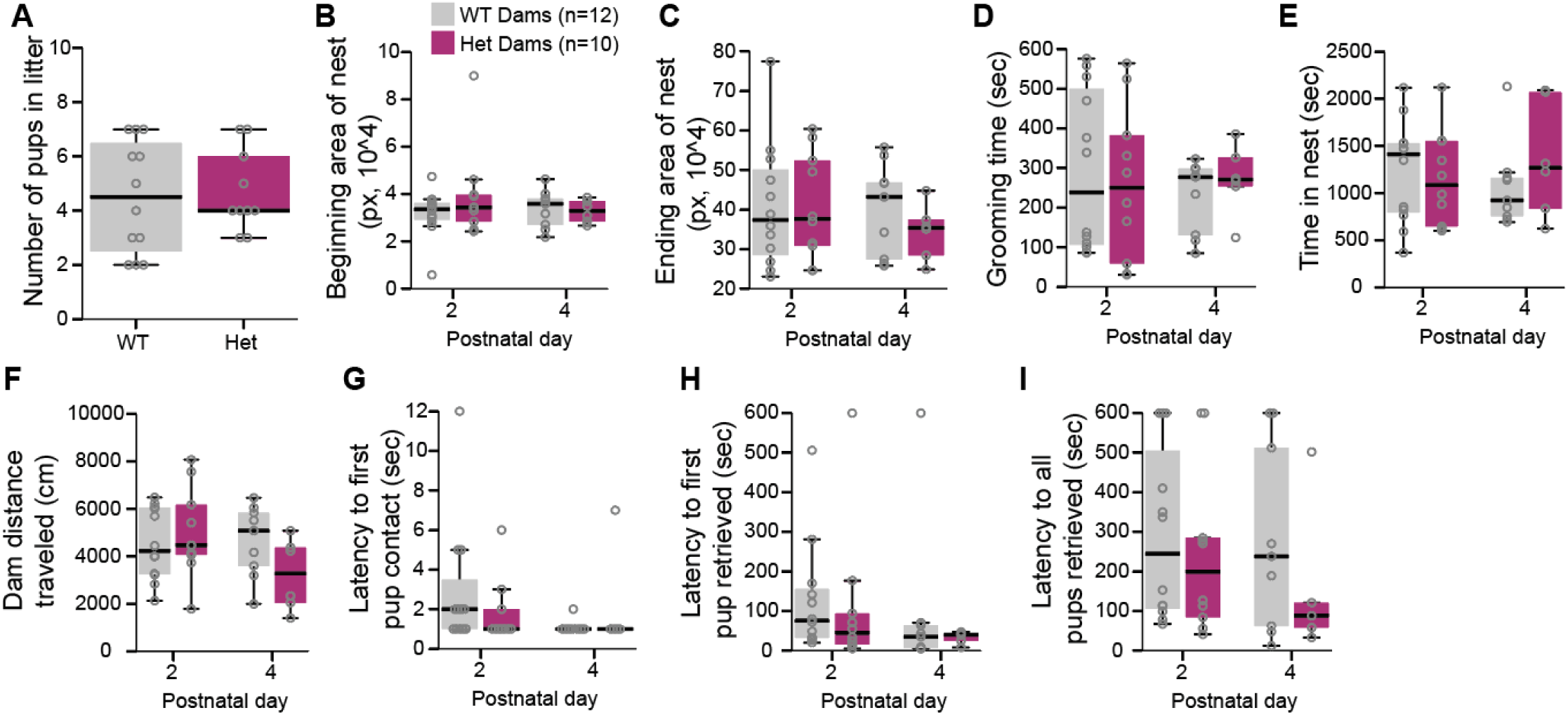
Maternal behavior was not altered by *Myt1l* mutation. (A) Litters produced by WT or Het dams were not different in number of pups. (B-C) The size of the nest area at the beginning (B) and end (C) of the maternal assessment session was not different between WT or Het dams at either postnatal pup age. (D-E) The time spent grooming (D) or in the nest (E) putatively attending to the pups were not different between genotypes at P2 or P4 assessment. (F) The WT and Het dams moved comparable distances around the cage during maternal care assessment. (G-I) During pup retrieval assessment, the WT and Het dams exhibited similar latencies to first contact (G), first retrieval (H), and last retrieval of the pups (I). Sample size of dams for all assessments is presented in panel B. Data are presented as boxplots respective group medians as, boxes 25^th^ – 75^th^ percentiles, and whiskers 1.5 x IQR. Individual data points are presented as open circles.

### MYT1L haploinsufficiency did not enhance agonistic behavior or alter home cages social hierarchies

Substantial evidence exists for the role of OXT in various social behaviors, which is supported by the fact that OXT projections and receptors are widespread through brain areas involved in social behavior (Dölen et al., 2013; Hung et al., 2017; Marlin et al., 2015; Modi and Sahin, 2019; Nardou et al., 2019; Peñagarikano et al., 2015; Sgritta et al., 2019; Tan et al., 2019; Walsh et al., 2018). As a social species, mice create social dominance hierarchies within their social groups. In laboratory mice, these hierarchies are established within the home cage environment in late adolescents. Exogenous application of OXT has been shown to increase the perception of dominance (Teed et al., 2019), suggesting a role for this peptide in social hierarchy behavior. Previously, we found that MYT1L haploinsufficient mice displayed submissive behavior when paired with a WT sex-matched, non-cagemate control (Chen et al., 2021). To understand if MYT1L haploinsufficiency and altered OXT cell density influenced the stability of home cage hierarchies, we again examined social hierarchy behavior in mice that were housed in sex- and genotype-matched groups from weaning. At P110, we conducted a round-robin style dominance assessment across three days using the tube test assay (**Figure 5A**). With the exception of one instance in one cage, all other cages of three mice showed transitive ranking on each test day (**Figure 5B, Figure S1**) (van den Berg et al., 2015). The exception was a cage of Het males that showed a non-transitive relationship where each mouse was submissive in at least one bout (**Figure 5C, Figure S1:** cage 6 day 1). While transitive rankings within a given day were observed for 47/48 comparisons, the rankings for the individual mice were only stable about 50% of the time across days for male and female WT mice, and male Het mice. In contrast, female Het mice were more likely to remain at the same ranking across days (**Figure 5D**). Rank only changed by one spot (e.g., a ranking of 1 never became a rank of 3 and vice versa). This suggests a general stability of the mice to remain in the dominant end or the submissive end of hierarchy (**Figure S1**), yet no impact of genotype on this stability.

**Figure 5.**
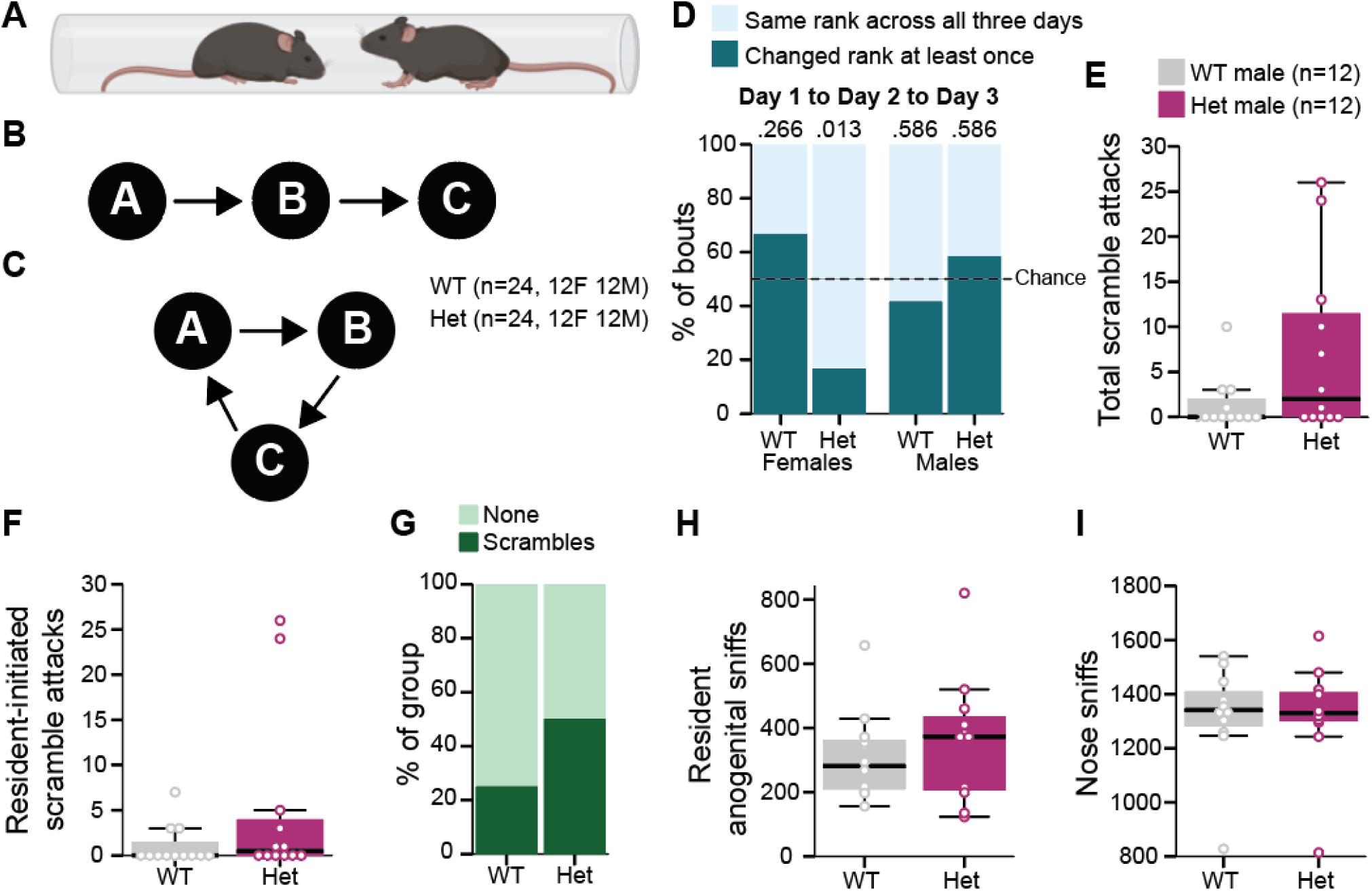
*Myt1l* mutation had a subtle, sex-specific effect on social rank change but did not influence male-specific aggression. (A) Schematic of the social dominance tube test assay. (B-C) Example of transitive (B) and non-transitive (C) dominance rankings within cages. (D) Only female Het mice showed significantly more instances of maintaining the same dominance rank across days compared to chance. WT females, WT males and Het males demonstrated chance levels for changing rank across days. Left panel: sample size for social dominance assessment. (E) In the resident intruder assay, Het and WT males did not differ significantly in the number of total scramble attacks that occurred. (F) Het and WT males initiated a significant number of scramble attacks. (G) *Myt1l* mutation was not associated with the occurrence of scramble attacks. (H-I) WT and Het males exhibited a similar number of anogenital sniffs (H) and nose-to-nose sniffs (I). Sample size of males for resident intruder paradigm is presented in panel E. Data in E,F, H & I are presented as boxplots respective group medians as, boxes 25^th^ – 75^th^ percentiles, and whiskers 1.5 x IQR. Individual data points are presented as open circles. Data in D & G are presented as counts.

Aggressiveness towards others has been reported in individuals with MYT1L syndrome (Coursimault et al., 2022). In addition, AVP activity in the hypothalamus plays a role in aggression in males (Terranova et al., 2017). Thus, to understand if MYT1L haploinsufficiency impacted agonistic and aggressive behavior, we assessed our male mice in the resident intruder paradigm. In response to a novel, age-matched intruder across three separate days, Het and WT resident mice engaged in and initiated a comparable number of scramble attacks with the intruder (**Figure 5E,F**). In addition, Het mutation was not associated with a difference in whether or not a scramble occurred at all (**Figure 5G**). Examination of non-aggressive social behaviors revealed similar frequencies of resident-initiated anogenital sniffs and nose-to-nose sniffs (**Figure 5H,I**) between Het and WT male littermates.

Thus, though OXT and AVP cell counts were abnormal, there did not appear to be an effect of this change on this set of hypothalamus-mediated behaviors. However, OXT was only one of several transcripts altered in the hypothalamus, and the downstream consequences of this multi-gene alteration are unknown. Therefore, we next conducted a broad survey of phenotypes regulated by the hypothalamus to determine other consequences of MYT1L heterozygosity on metabolic and behavioral traits.

### MYT1L haploinsufficiency had no effect on sexual maturity or endocrine organ weights

Endocrine issues have been reported in individuals with MYT1L Neurodevelopmental Syndrome, including early onset puberty (Windheuser et al., 2020). Thus, to examine the potential impact of altered neuropeptide expression within the hypothalamus on downstream hormonal axes and glands, we examined gonadal and adrenal weights of female and male Hets and WT littermates, as well as pubertal or sexual maturity onset. MYT1L genotype had no effect on combined adrenal weights in either females or males (**Figure 6A**). Hets and WTs had no difference in gonadal weights, including ovaries or testes (**Figure 6B,C**), nor did these individuals differ in overall body weight (**Figure 6D**). Next, we examined preputial separation and vaginal openings in young male and female, respectively, Het and WT littermates. No differences in age at reaching these anatomical markers of sexual maturity were observed (**Figure 6E**), and the mice were of equal weight when this happened (**Figure 6F**). Thus, MYT1L haploinsufficiency did not affect size of gonads or onset of sexual maturity.

**Figure 6.**
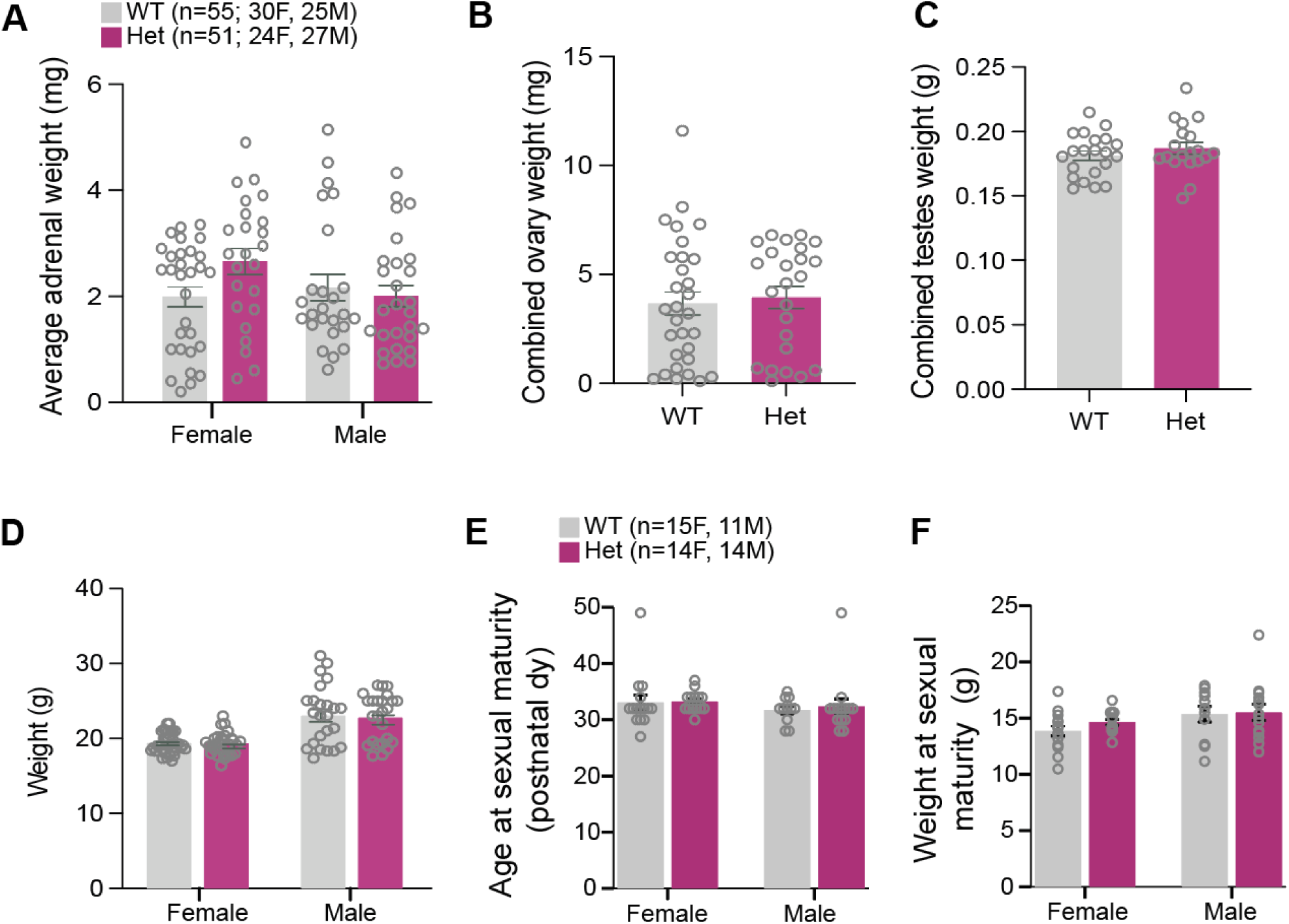
MYT1L haploinsufficiency showed no effect on sexual maturity. (A) Female and male Het mice have similar average adrenal weights compared to WTs. (B-C) Combined ovary weights in female Het mice (B) and combined testes weights in male Het mice (C) were comparable to WTs. (D) Body weights of these mice were also similar between genotypes. (E-F) WT and Het female and male littermates reached sexual maturity at a similar age (E) and at a similar weight (F). Sample size for gland weights is presented in panel A. Sample size for sexual maturity assessment is presented in panel E. Data are represented as means ± SEM with individual points presented as open circles.

### MYT1L haploinsufficiency alone did not induce metabolic disruption or persistent feeding changes

*MYT1L* loss-of-function is associated with childhood obesity, however, with variable reporting of parallel hyperphagia (Blanchet et al., 2017; Carvalho et al., 2021; Coursimault et al., 2022). Thus, loss of MYT1L function may alter motivation to consume food leading to increased weight gain and obesity. Alternatively, increased weight gain and resulting obesity may be due to an underlying metabolic dysfunction that is either independent from or precedes aberrant feeding behavior. Previously, we observed increased weight in Het mice with an onset in early adulthood, which persisted throughout life for that specific cohort (Chen et al., 2021; Schreiber et al., 2024). To understand if MTY1L haploinsufficiency results in aberrant metabolic or behavioral processing that may drive differences in weight, we examined metabolic markers as well as feeding behavior and motivation to obtain and consume food both at a young juvenile age and again in adulthood (**Figure 7A**).

**Figure 7.**
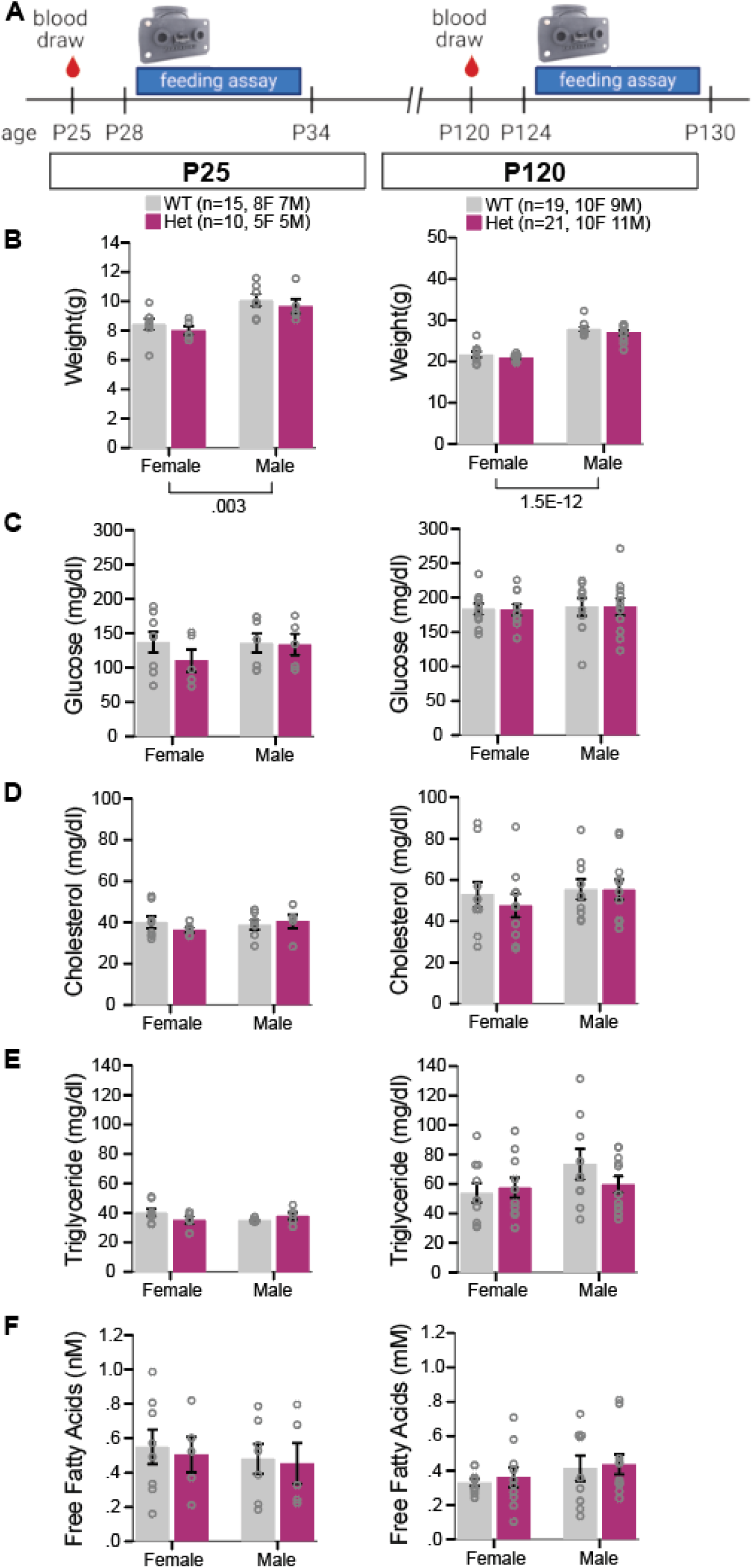
*Myt1l* mutation did not impact weight or metabolic markers in juvenile or early adult mice. (A) Schematic of experimental timeline that included blood draw for metabolic marker assessment and feeding assay at two ages. (B) Het female and male mice weighed comparably to WT littermates on regular chow at both P25 (left panel) and P120 (right panel). (C-F) Het mice did not differ in levels of fasted glucose (C), cholesterol (D), triglycerides (E) or free fatty acids (F) from WT littermates at P25 (left panels) or P120 (right panels). Sample sizes for weight and metabolic assessments are presented in panel B. Data are represented as means ± SEM with individual points presented as open circles.

Markers of metabolic disruptions including those of glucose, cholesterol, triglycerides, and free fatty acids may precede onset of weight gain or result from it. To understand if MYT1L haploinsufficiency results in metabolic dysfunction, we examined these metabolic markers in juvenile and adult mice. In this cohort, at blood draw, no differences in weight were observed between Hets and WT littermates at either age point, although males weighed significantly more than females at both ages, as expected (**Figure 7B**). No differences were observed in fasted blood glucose, cholesterol, triglycerides, or free fatty acids at either time point (**Figure 7C-F**), indicating in the absence of greater weight, MTY1L haploinsufficiency did not result in metabolic dysfunction.

To test feeding motivation, we used programmable feeding devices to monitor feeding behavior across daily cycles and assessed how hard Het and WT mice would “work” to obtain food pellets via an operant conditioning paradigm. The feeding was assessed in four phases: magazine training, dual active, fixed ratio 1 (FR1), and resetting progressive ratio (**Figure 8A**). During magazine training, when the device consistently offered a food pellet without requiring nose pokes, the Het mice removed more freely available pellets than WT mice during the inactive phase, regardless of age (**Figure 8B**). In contrast, when the mice were required to start working for the pellets during the Dual Active phase, when both nose poke holes responded with a pellet, each group achieved a comparable number of pellets (**Figure 8C**) by engaging in a comparable number of pokes across both the active and inactive phases (**Figure 8D**). During FR1 training, when only one hole produced a pellet, both Hets and WTs learned to nosepoke in the correct hole for access to pellets (**Figure 8E**), with no difference in the number of pellets achieved (**Figure 8F**).

**Figure 8.**
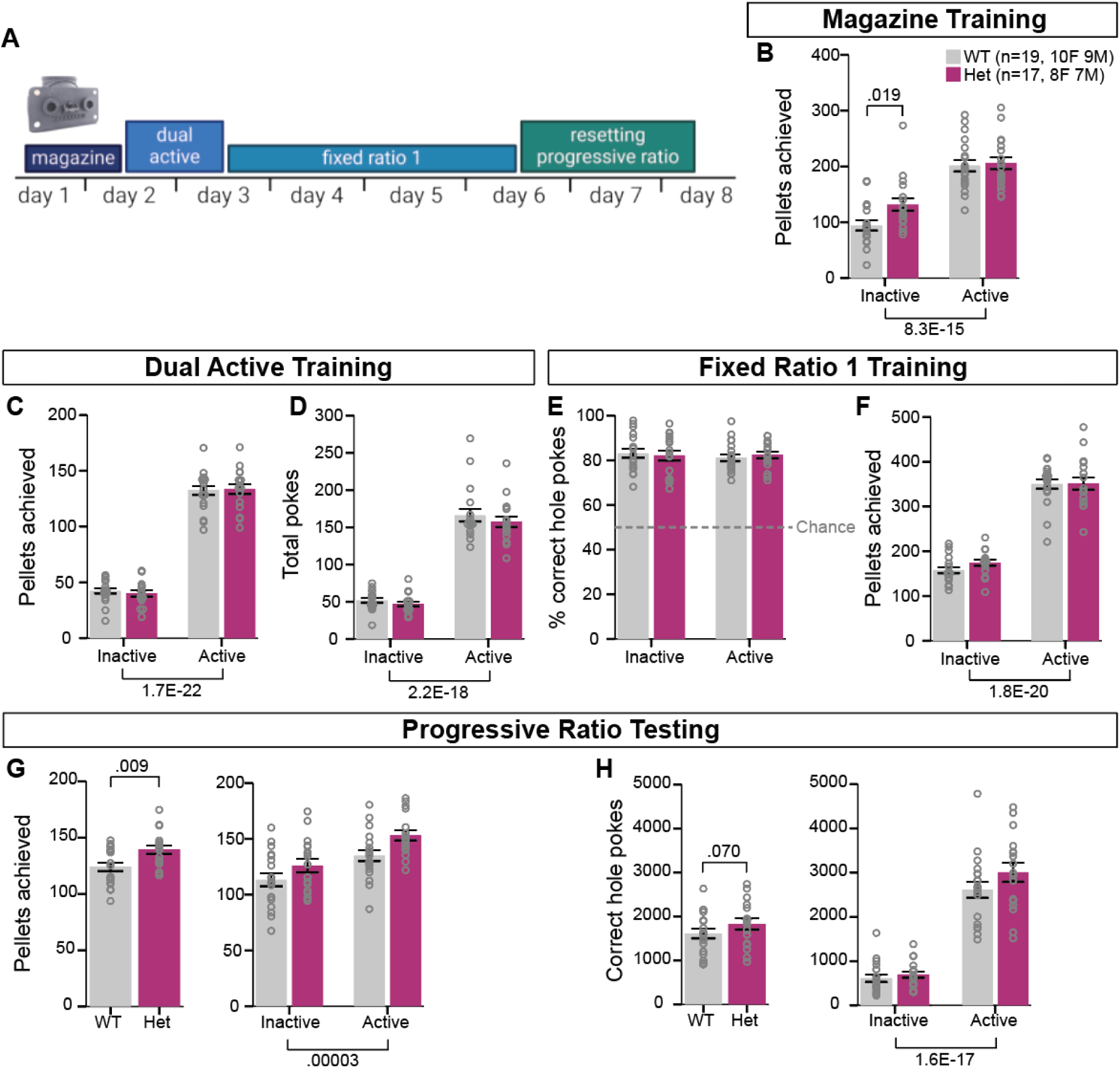
*Myt1l* mutation impacted feeding in a high effort situation. (A) Schematic of the feeding assay timeline. (B) During magazine training, Het mice consumed more pellets during only the inactive phase than WT littermates. (C-D) During dual active training, Het and WT mice consumed a comparable number of pellets (C) and exhibited a similar number of nose pokes (D). (E-F) During FR1 training, Het mice poked into the correct hole at a comparable percent to WT mice (E), resulting in a similar number of pellets consumed between groups (F). (G-H) During resetting PR testing, Het mice consumed more pellets (G) and exhibited a marginal increase in correct hole pokes (H). Sample size for the feeding assay in panel B. Data are represented as means ± SEM with individual points presented as open circles.

Finally, the mice were tested in the resetting progressive ratio phase, which requires an increasing number of pokes to achieve each subsequent pellet. A trial would end and reset back to 1 poke required if 30 minutes passed of inactivity. By increasing the effort required for subsequent pellets we are able to better quantify the motivation of the animals to acquire pellets. The Het mice achieved a greater number of pellets compared to WT mice (**Figure 8G**), with a non-significant increase in correct hole pokes (**Figure 8H**), suggesting an increase in engagement with the PR task.

The increase in pellets achieved during the PR task in regular chow-fed Het mice suggests something about the variable reward value induced a higher engagement with the task. It is unlikely that the task induced them to eat more, as mice typically eat a consistent amount of food across days (Matikainen-Ankney et al., 2021; Nguyen et al., 2016), and an increase in pellets was not observed across tasks. The slight increase in correct hole pokes suggests a potential perseveration in the Hets to continue to engage with the task and a resistance to extinction at high efforts. However, overall, the Het and WT mice leveraged a similar strategy in this task - long trials of high hole poke effort - to achieve pellets. All together, these data indicate that, in the absence of enhanced weight gain, MTY1L haploinsufficiency did not affect a subset of markers of metabolic health yet subtly impacted feeding behavior.

### MYT1L haploinsufficiency interacted with diet to influence weight

In our initial characterization of MYT1L haploinsufficiency in the mouse, increased weight in the Het mice on a regular chow diet was observable in adulthood by P94 (Chen et al., 2021). However, in the current study, we did not observe a distinction between the weight of Het and WT littermates on a regular chow diet by P120 in males or females (**Figures 7B, S2A**).

However, not all individuals with a putative *MTY1L* loss-of-function variant become obese (Blanchet et al., 2017; Carvalho et al., 2021; Coursimault et al., 2022). Thus, *MTY1L* mutations are an incompletely penetrant risk factor for childhood obesity. There may be other risk factors that interact with this genetic vulnerability to drive in this phenotype. To probe the possibility that diet interacts with MTY1L variation, we fed our mice a high fat diet (HFD) and again examined weight, metabolic markers, and feeding behavior. Genotype-matched cages of 3-5 Hets or WT littermates were given access to either regular chow or HFD for 14 weeks starting at P21. Both male and female Het mice gained significantly more weight on HFD than their WT littermates (**Figure 9A, S2B**), indicating that depending on the diet, Het mice will gain more weight.

**Figure 9.**
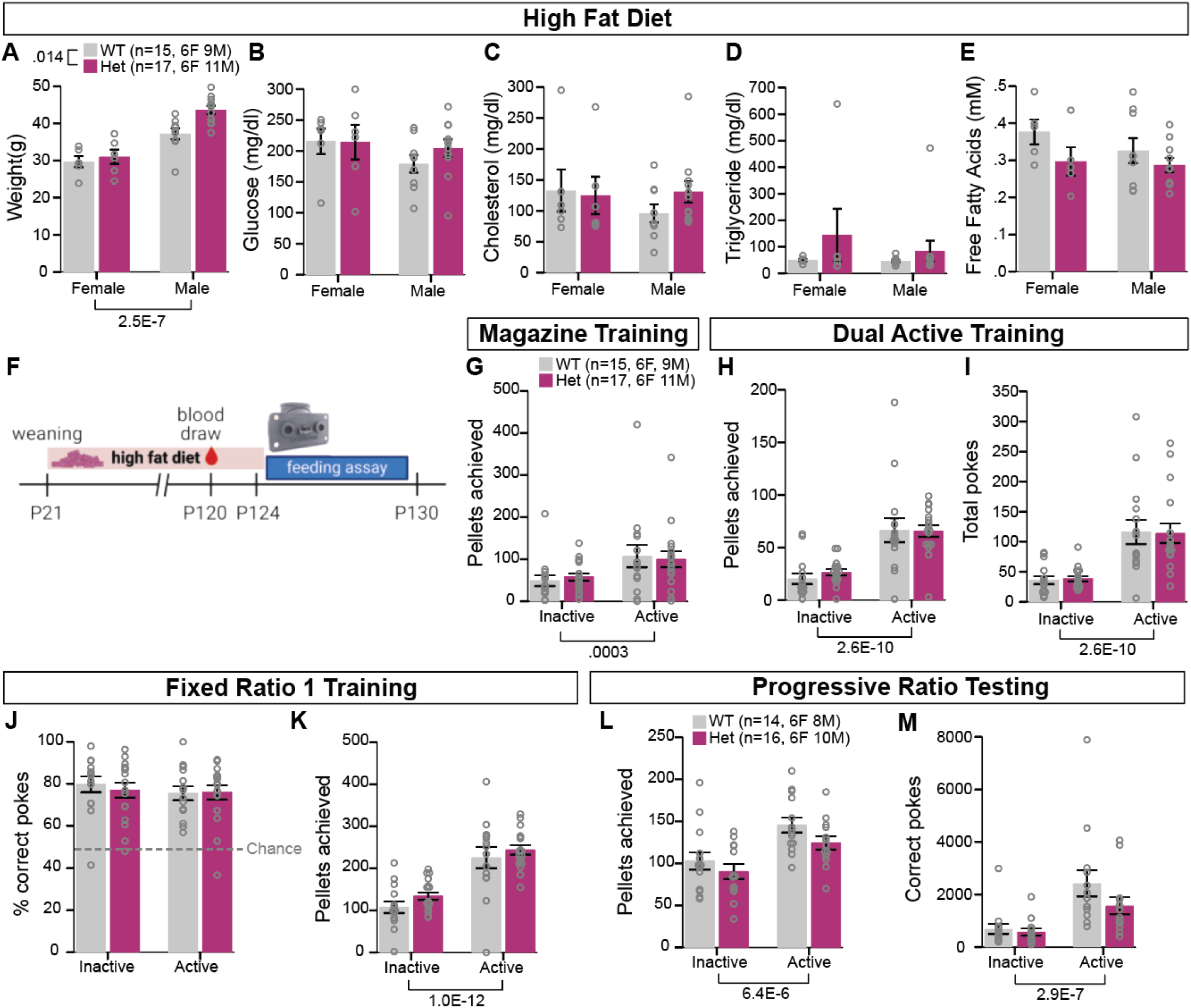
HFD interacted with *Myt1l* mutation to influence weight with unchanged metabolic markers and feeding performance. (A) Following 14 weeks of HFD, Het mice weighed significantly more than WT littermates at P120. (B-E) HFD-fed Het mice did not differ in levels of fasted glucose (B), cholesterol (C), triglycerides (D) or free fatty acids (E) from HFD-fed WT mice. (F) Schematic of experimental timeline that included blood draw for metabolic marker assessment and feeding assay following 14 weeks of HFD. (G) During magazine training, Het and WT mice consumed a comparable number of pellets. (H-I) During dual active training, Het and WT mice consumed a comparable number of pellets (H) and exhibited a similar number of nose pokes (I). (J-K) During FR1 training, Het mice poked into the correct hole at a comparable rate to WT mice (I), resulting in a similar number of pellets consumed between groups (K). (L-M) During resetting PR testing, Het mice consumed a comparable number of pellets as WT mice (L) and exhibited a comparable number of correct pokes (M). Sample size for metabolic marker assessment, magazine, dual active and FR1 feeding assessments in panel A. Sample size for PR testing in panel L. Data are represented as means ± SEM with individual points presented as open circles.

Possible changes in metabolic function were then tested at P120 following the HFD-induced weight gain. Despite the increased weight at blood draw (**Figure 19A**), all markers tested were comparable in Het mice to WT littermates (**Figure 9B-E**). This indicates that there were no gross metabolic anomalies as a consequence of MTY1L haploinsufficiency and is consistent with the increased weight being more likely related to altered feeding behavior.

To test feeding behavior and motivation across the circadian cycle, the mice fed HFD for 14 weeks were assessed in our operant feeding paradigm (**Figure 9F**). No differences between groups were observed in pellets achieved during the free feeding magazine training (**Figure 9G**) or dual active training (**Figure 9H**), with a comparable number of total pokes as well (**Figure 9I**). During FR1 training, both Hets and WTs learned to nosepoke in the correct hole to receive a pellet with comparable rate (**Figure 9J**), achieving a similar number of pellets (**Figure 9K**). Finally, during the resetting PR task, HFD did not impact motivation to feed, demonstrated by comparable pellets achieved (**Figure 9L**) and correct hole pokes for Hets and WTs (**Figure 9M**). Thus, while HFD differentially impacted weight in the Het mice, this did not influence feeding behavior in this task.

## Discussion

MYT1L haploinsufficiency in the mouse recapitulates many clinically-relevant phenotypes (Chen et al., 2021; Kim et al., 2022; Wöhr et al., 2022) providing a model to aid our efforts to better understand brain region and cell type specific changes that might be targeted for interventions. In the current study, we leveraged one such model to investigate the participation of MYT1L in hypothalamic and hypothalamus-mediated phenotypes. The hypothalamus is of particular interest here based on patient reports of disruptions to endocrine-related processes (Coursimault et al., 2022) and previous work in other experimental systems showing cell changes to hypothalamic brain areas in the absence of *Myt1l* orthologs (Blanchet et al., 2017). Here, we uncovered epigenetic and transcriptomic changes suggestive of context- and region-specific roles for MYT1L, including regulation of neuropeptide systems. Behaviorally, we found the MYT1L haploinsufficient mouse largely unchanged in hypothalamus-targeted phenotypes.

As in previous cortex studies, we see that MYT1L loss results in altered chromatin state and gene expression in the hypothalamus. In the adult murine cortex, approximately 10,000 DARs were identified with ATAC-seq (Chen et al., 2021). Here, in the hypothalamus, we see that MYT1L haploinsufficiency results in almost 13,000 DARs. Approximately 3,000 of these regions show up as differentially accessible in both cortex and hypothalamus datasets. So, while there do exist shared consequences of MYT1L haploinsufficiency on chromatin accessibility in both brain regions, there are clearly numerous region-specific effects as well. In concordance with cortex studies, we see TF motifs in more accessible promoter regions with functions relating to overall neurodevelopment, along with more specific functions including differentiation and neuronal activity (e.g., ETS1, THAP11, ELF4, ZFX, and NFYA/B). Interestingly, we also see enrichment in the KLF14 TF motif in more accessible regions with MYT1L haploinsufficiency. KLF14 is known to be a master regulator of various metabolism related pathways (Akash et al., 2023). In enhancer regions, we still see that more accessible regions contain TF motifs related to neuronal differentiation (e.g., RFX2, DLX2, NEUROD1, CTCF) in concordance with cortex studies. However, we also see that in the hypothalamus MYT1L haploinsufficiency leads to more accessible regions with TF motif enrichment related to activity dependent genes (e.g., MEF2A, JDP2, JUND). Regarding differential gene expression, previous work in cortex identified 533 DEGs and here in hypothalamus we see 718 DEGs with MYT1L haploinsufficiency (Chen et al., 2021). Only 44 genes are differentially expressed in both cortex and hypothalamus, suggesting that MYT1L may confer different effects on gene expression between brain regions. While specific genes may not show an overlap, we do see a general agreement between the two regions in that MYT1L loss leads to upregulation of genes involved in early neuronal differentiation. Finally, we see that MYT1L haploinsufficiency seems to activate genes involved in neuropeptide hormone expression in the hypothalamus.

We observed a decrease in AVP positive cell numbers in the PVN of the hypothalamus in MYT1L haploinsufficiency. This replicates previous work in zebrafish in which MYT1L knockdown resulted in decreased AVP expression in the neuroendocrine hypothalamus (Blanchet et al. 2017), an area that functions similarly to the mammalian PVN. Our results, however, did not replicate the previously reported decrease in OXT expression also observed in the zebrafish neuroendocrine hypothalamus (Blanchet et al., 2017). This may reflect subtle differences between models or methods. Unlike Blanchet et. al., which conducted a full morpholino knockdown, clinically-relevant haploinsufficiency modeled here might not fully recapitulate all the molecular consequences of complete loss during hypothalamic development. Furthermore, the previous publication used *in situ* hybridization for mRNA localization, whereas we examined OXT protein changes at the level of the cell. Thus, MYT1L loss may disrupt OXT at the RNA level which might fail to translate to changes in cell number expression of the protein. One way to clarify this is single cell/nuclei RNA-sequencing in the haploinsufficient brain to directly examine cell proportion and RNA expression at the cell type-specific level.

Our findings that MYT1L loss leads to disrupted transcription of neuropeptide regulation in the adult hypothalamus and changed OXT/AVP-positive cell numbers did not line up with consequences on behaviors known to respond to neuropeptide activity. Specifically, maternal behaviors and aggression, two social domains shaped by OXT and AVP, respectively (Champagne et al., 2001; Insel et al., 1997; Pedersen et al., 1982; Pedersen and Prange, 1979; Terranova et al., 2017), were unchanged. The C57BL/6J mouse line, on which the *Myt1l* Het line was generated, is not a highly aggressive strain of mice (Lidster et al., 2019; Weber et al., 2022). Thus, the low number of scramble attacks observed overall is not unexpected. Regardless, the Het males had a non-significant increase in aggressive displays and percentage of Het males with an aggressive display at all compared to WT males, suggesting a potentially smaller effect than we were able to detect. Follow up studies with greater power and in a more aggressive strain, such as CD1, may yield further information on the impact of *Myt1l* mutation on agnostic and aggressive behaviors.

The lack of weight gain by Hets fed regular chow in this study contrasts with our previous work with this same model and others (Chen et al., 2021; Wöhr et al., 2022). This could be driven by several different factors. The effect of MYT1L haploinsufficiency on weight could be smaller than previously concluded, with variability in how the phenotype expresses. In humans, while MYT1L mutation is a risk for childhood obesity (Carvalho et al., 2021; Kalinderi et al., 2024), this is incompletely penetrant in the population, with the largest study to date reporting a 58% rate of obesity and overweight (Coursimault et al., 2022). This incomplete penetrance suggests other interacting factors. One such factor could be diet, which is supported by our data here. Variability may also interact with age. Our previous report observed a significant weight gain by P94 (Chen et al., 2021), yet none by P120 here. Other genetic models of MYT1L haploinsufficiency revealed significant weight increase only after P120 (Wöhr et al., 2022) or a trend as late as P236 (Kim et al., 2022). Yet another study did not report weight phenotype at all, although the mice were assessed only until two months of age (Weigel et al., 2023). In the two previous reports of significant weight gain, the effects were largest in females and more variable than WTs. Finally, age-related severity in other phenotypic consequences of MYT1L loss has been shown (Kim et al., 2022). It will be important to evaluate this phenotypic variation further in subsequent cohorts with greater power for sex by genotype interactions and at later ages.

This study further defined a lack of impact of MYT1L haploinsufficiency alone, as modeled here, on markers of gross metabolic health and feeding motivation. HFD interacted with MYT1L haploinsufficiency to increase weight, yet we could not define a metabolic change, within our measurements, or enhanced feeding behavior that paralleled this phenotype. However, our investigation was not exhaustive and weight gain is a complex process. Further dissection of the HFD-induced weight phenotype is warranted in future studies. Recent work with a similar paradigm employed here revealed that requiring even small effort - one nose poke - to achieve food pellets decreases food intake even in models of diet-induced obesity (Barrett et al., 2024). Thus, the effort required by our task may have masked any increase in food intake induced by the mutation. This is supported by the increase in pellets taken by Het mice during the first 24 hours of the task, when pellets were freely available.

While an overall increase in feeding motivation was not revealed by the feeding operant experiments here, regular chow-fed MYT1L mice increased engagement with the task in the form of increased pellets achieved during the progressive ratio portion. Previously, we observed a reduction in social interactions achieved by male Hets in a social operant paradigm (Chen et al., 2021). Together, these data suggest MYT1L haploinsufficiency reduces interest in working for social interactions but may increase interest in working for food, although this is confounded by potential overall decrease in food intake when effort is required. Future studies directly comparing operant-based consumptive behavior for social interaction rewards to that for highly palatable food rewards (e.g., sucrose pellets) will be important for clarifying these phenotypes. HFD has been shown to reduce motivation even for sweet palatable foods (Arcego et al., 2020), which may underlie the lack of similar increase in the HFD-fed Het mice. More work needs to be done to illuminate the interaction between HFD and *Myt1l* mutation on motivational behaviors.

It is notable that in humans, MYT1L mutations result in a highly variable phenotype. While developmental and behavioral disorders are observed in most individuals, extra-neurological features are less common. For example, endocrine and lipid abnormalities have been reported in only 3% to 18% of individuals (Coursimault et al., 2022), including a variety of conditions such as hypoprolactinemia, hypothyroidism, hypogonadism, and dyslipidemia. Yet, these rates are much higher than the general population. This suggests that MYT1L haploinsufficiency alone is insufficient to cause these abnormalities, but that it may interact with other risk factors, environmental or genetic, to drive these specific phenomena. This suggests that crossing our models to different genetic backgrounds, or exposing them to more environmental risk factors (as hinted at in our high-fat diet experiments), could unmask the mechanisms by which this occurs.

## Conclusions

Understanding the impact of MTY1L loss on specific brain regions and cell-types will enable more refined mechanistic and interventional studies that may lead to targeted therapies for MYT1L Neurodevelopmental Syndrome. We have shown that MYT1L haploinsufficiency results in epigenetic and transcriptomic consequences in the adult hypothalamus that influence neuropeptide systems and changes in cell distributions. These data suggest targeting these systems may provide useful therapeutic avenues. However, we did not observe altered typical sex differentiated behaviors, specifically maternal care and territorial aggression, nor significantly increased motivation to obtain food - behaviors with a known hypothalamic role. Nor were signatures of gross endocrine dysfunction altered, suggesting that these autonomic systems are largely intact in the context of MYT1L haploinsufficiency. Yet, these are not all the phenotypes or processes driven by this so-called endocrine system master gland, and further examinations are necessary. Our findings, therefore, do not exclude the hypothalamus as an important area in the etiology of MYT1L Neurodevelopmental Syndrome, but do indicate the specific behaviors assessed here are largely normal. It is quite possible the epigenetic and transcriptomic changes in this area have greater impacts on circuit function elsewhere, as systems like the oxytocinergic have widespread projections. Nevertheless, investigations into the role of MYT1L in other brain regions and cell types to understand robust phenotypes such as ADHD-related hyperactivity will be important next steps.

## Acknowledgements

This work was supported by The Jakob Gene Fund, SFARI, McDonnell International Scholars Academy (to JC), and the National Institutes of Health: R01MH124808 to KLK, JDD & SEM, P30DK56341 to WUSTL Nutrition Obesity Research Center (NORC) and P50HD103525 to IDDRC@WUSTL. We thank Dr. Alexxai Kravtiz for valuable discussions, Dr. Sangeeta Adak of the NORC Animal Research Core for her assistance with metabolic assessments, Ms. Kelly Kim for frame labeling assistance, and the McDonnell Genome Institute for sequencing services.

## Author contributions

Conceptualization, SEM, JC, JDD; Data curation, SEM, KBM, SMC, DS; Formal analysis, SEM, SMC, DS, JC, DW; Funding acquisition, SEM, KLK, JDD; Investigation, KBM, SMC, DS, JC; Methodology, SEM, KBM; Project administration, SEM, JDD; Resources, SEM; Software, KBM; Supervision, SEM, JDD; Visualization, SEM, SMC, DS, DW, RC; Writing - original draft, SEM, KMB, SMC, DS, DW, JLG, RC; and Writing - review & editing, all authors.

## Declaration of Generative AI and AI-assisted technologies in the writing process

None

## Supplementary Material

**Supplementary Figure 1.**
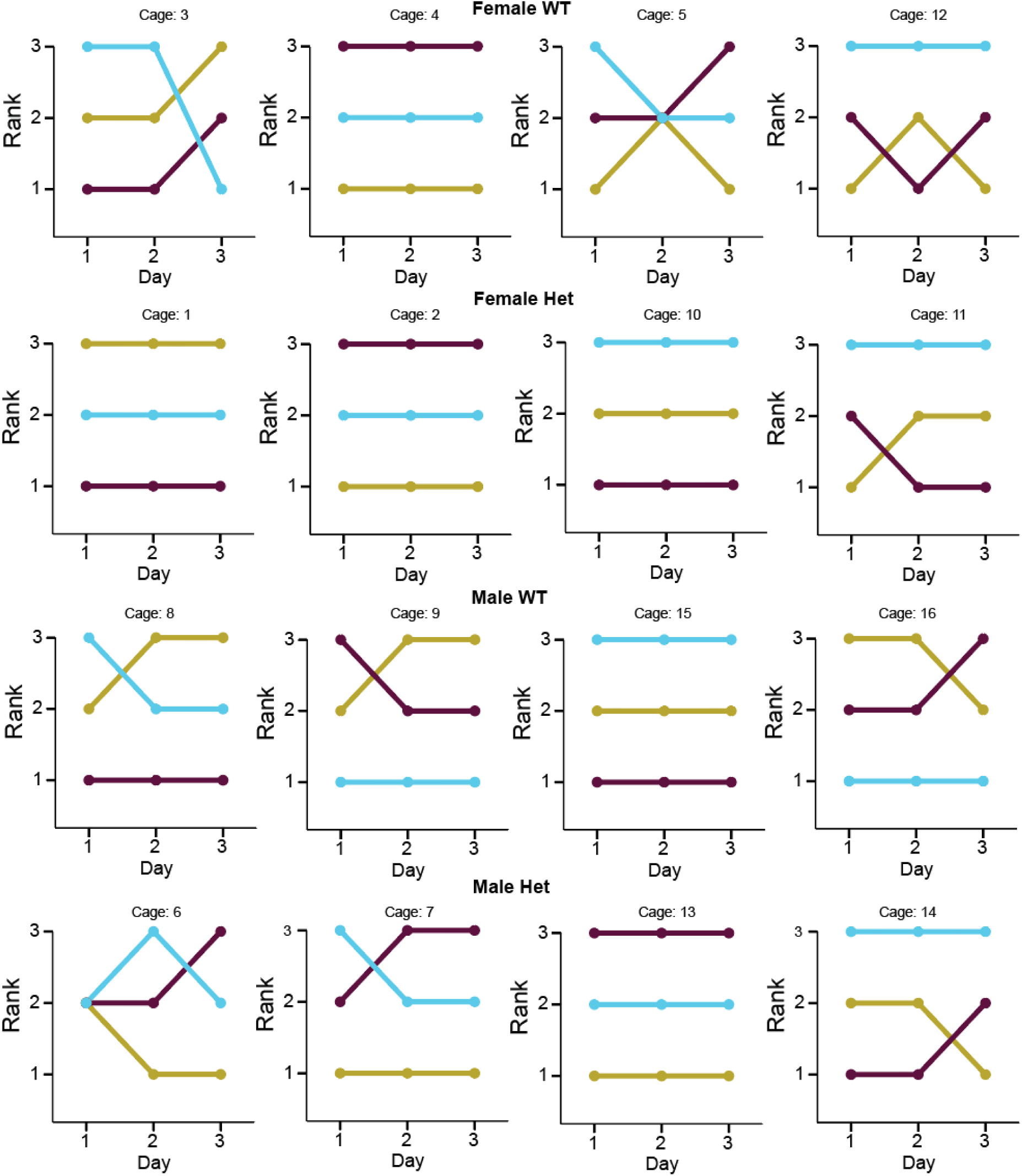
Daily rankings for male and female WT and Het cages in the three day, round robin social hierarchy assessment.

**Supplementary Figure 2.**
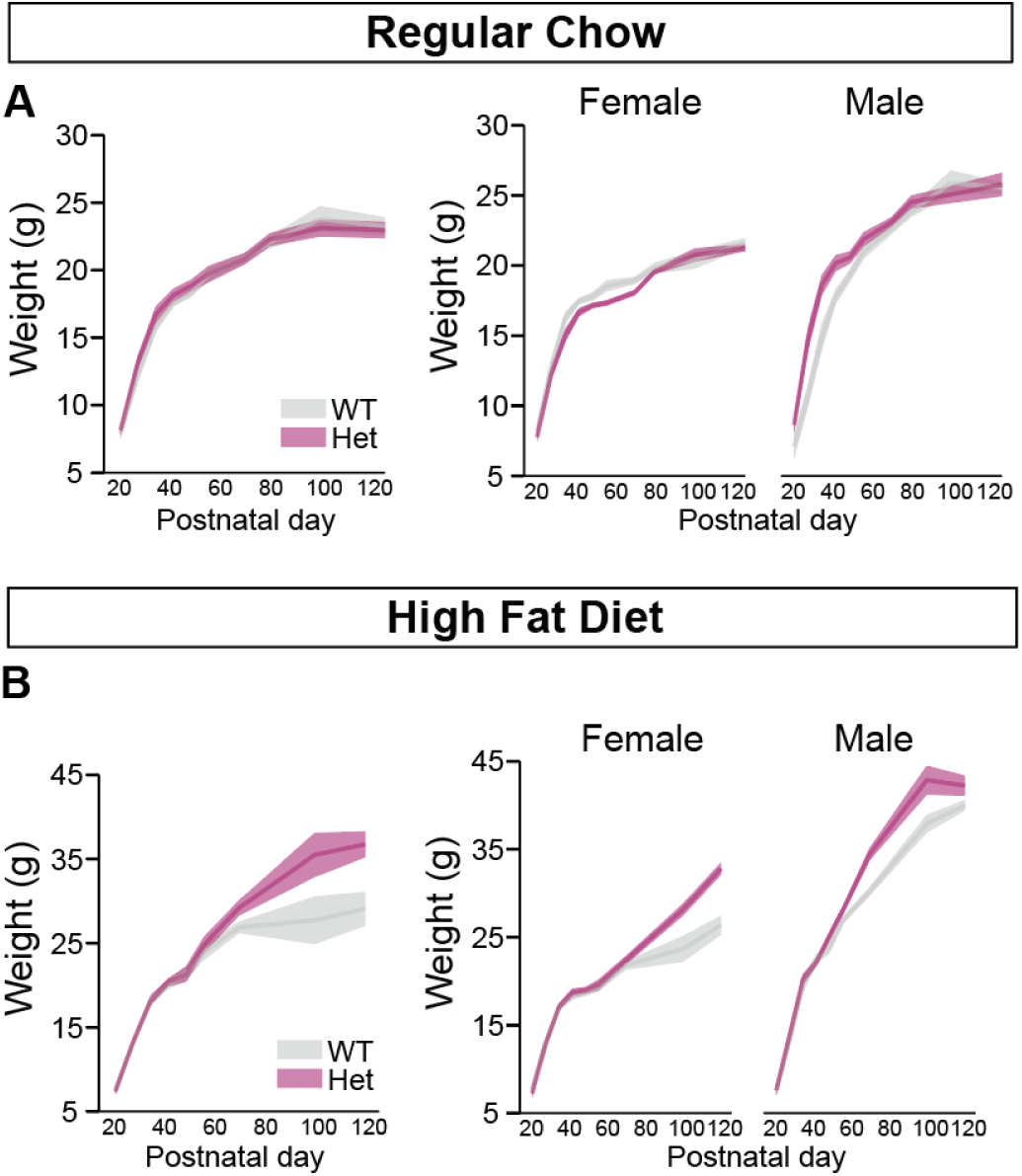
*Myt1l* mutation interacted with a high fat diet to impact weight. (A) When fed regular chow, Het females and males gained weight at a comparable rate to WT littermates until early adulthood (P120). (B) When fed HFD for 14 weeks, Het females and males gained weight at a significantly greater rate than WT littermates. Data are represented as means ± SEM.

**Supplementary Table 1. Differential epigenetic and transcriptomic analyses of the MYT1L haploinsufficient hypothalamus.**

