## Supplementary Material for "A survey of hypothalamic phenotypes identifies molecular and behavioral consequences of MYT1L haploinsufficiency in male and female mice"

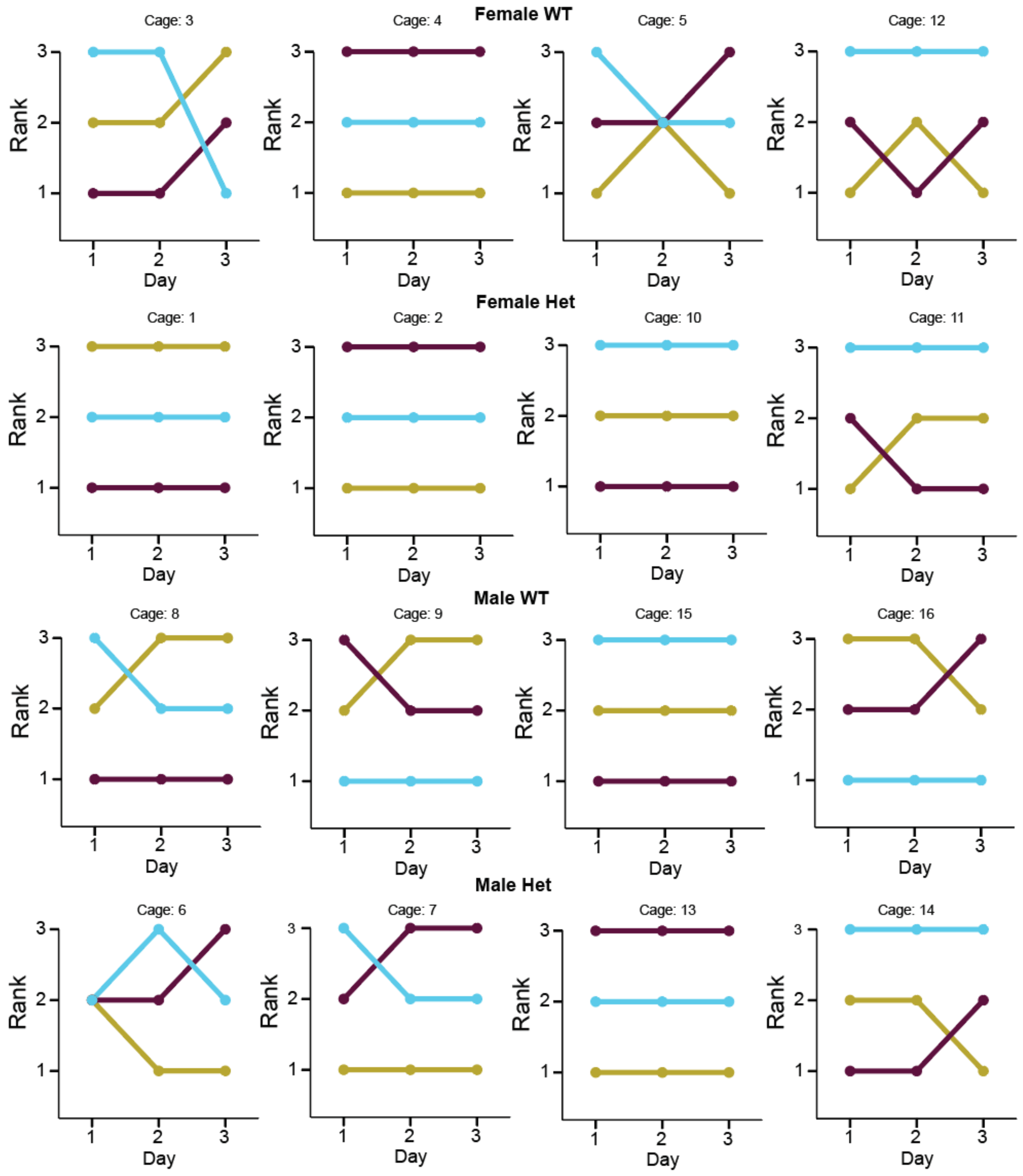


**Supplementary Figure 1. Daily rankings for male and female WT and Het cages in the three day, round robin social hierarchy assessment.**

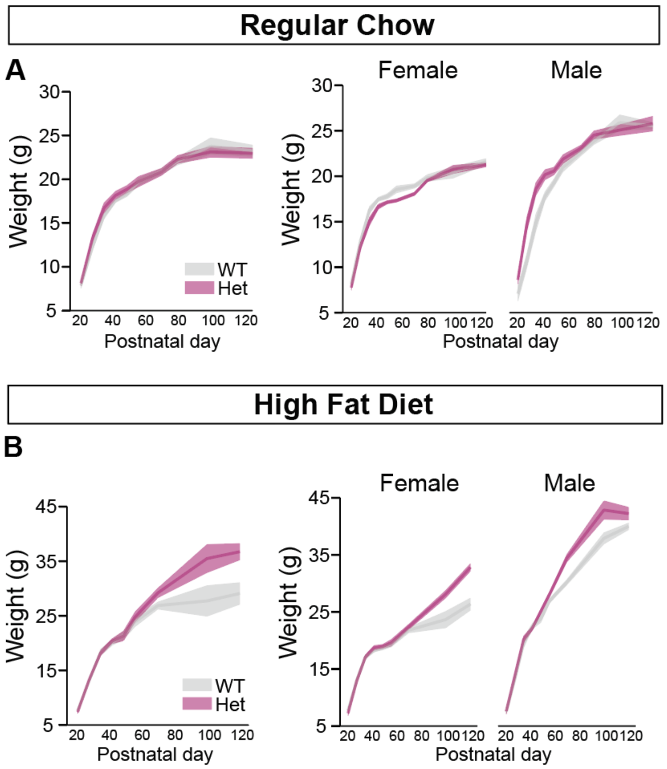


**Supplementary Figure 2.** ***Myt1l*** **mutation interacted with a high fat diet to impact weight.** (A) When fed regular chow, Het females and males gained weight at a comparable rate to WT littermates until early adulthood (P120). (B) When fed HFD for 14 weeks, Het females and males gained weight at a significantly greater rate than WT littermates. Data are represented as means ± SEM.

**Supplementary Table 1. Differential epigenetic and transcriptomic analyses of the MYT1L haploinsufficient hypothalamus.**
